# Riding the Wave: Unveiling the Conformational Waves from RBD to ACE2

**DOI:** 10.1101/2023.05.12.540230

**Authors:** Nikhil Maroli

## Abstract

The binding affinity between angiotensin-converting enzyme 2 (ACE2) and the receptor-binding domain (RBD) plays a crucial role in the transmission and re-infection of SARS-CoV2. Here, microsecond molecular dynamics simulations revealed that point mutations in the RBD domain induced conformational transitions that determined the binding affinity between ACE2 and RBD. These structural changes propagate through the RBD domain, altering the orientation of both ACE2 and RBD residues at the binding site. ACE2 receptor shows significant structural heterogeneity, whereas its binding to the RBD domain indicates a much greater degree of structural homogeneity. The receptor was more flexible in its unbound state, with the binding of RBD domains inducing structural transitions. The structural heterogeneity observed in the ACE2 unbound form plays a role in the promiscuity of viral entry as it may allow the receptor to interact with various related and unrelated ligands. Furthermore, rigidity may be important for stabilizing the complex and ensuring the proper orientation of the RBD-binding interface with ACE2. The greater structural homogeneity observed in the ACE2-RBD complex revealed the effectiveness of neutralizing antibodies and vaccines that are primarily directed towards the RBD-binding interface. The binding of the B38 monoclonal antibody revealed restricted conformational transitions in the RBD and ACE2 receptor, attributed to its potent binding interaction.

## Introduction

The SARS-CoV2 is a positive-strain single-stranded RNA virus that is causing the ongoing coronavirus disease 2019 (COVID-19) pandemic. Its highly transmissible and pathogenic properties have led to high morbidity and mortality rates worldwide since its initial detection in December 2019^1^. Earlier studies have demonstrated the crucial involvement of angiotensin-converting enzyme 2 receptor in facilitating viral infections ^2, 3^. The entry of viral particles is mediated by angiotensin-converting enzyme 2, which is expressed on the surface of various organs of human cells. Binding of the viral spike glycoprotein and ACE2 determines transmissibility and replication in host cells^4–7^. The S protein initiates receptor binding to the S1 and S2 subunits, which then fuses and initiates attachment to the host membrane^8^. The primary role of ACE2 is to hydrolyze angiotensin 2, thereby lowering blood pressure^9^. The receptor is mainly expressed in the alveolar epithelial cells of the lungs. To achieve a higher affinity for spike protein binding, ACE2 must maintain a receptor-accessible state with at least one upward conformation to avoid steric clashes that disrupt the binding process^10, 11^. SARS-CoV2 is rapidly spreading worldwide, with several new variants exhibiting stronger affinity for its receptors^12^. The lineage B.1.1.7 or N501Y.V1 was first detected in England and rapidly expanded to over 25 countries^13–15^. Lineage B.1.351 or N501Y.V2 was discovered in South Africa and was responsible for most pandemics at the end of 2020^16, 17^. Experimental and theoretical studies have revealed that mutations result in a higher binding affinity for ACE2-RBD^18–21^. Furthermore, the mutations E484K and K417N showed a higher binding affinity between the RBD and ACE2^19^. Lineages B.1.617.2, G/452R. V3, 21A, and 21A/S: 478 K have been reported in more than 100 countries. In 2021, the emergence of lineage B.1.1.529, which contains multiple mutations in spike proteins, resulted in increased transmissibility, immune system evasion, and vaccine resistance^21–24^. However, these mutations have been shown to cause fewer severe illnesses than in previously reported strains. It has been shown that antibody serum-based therapy may aid in the recovery of patients infected with the virus^25^.

Binding of the S protein to ACE2 causes conformational changes that led to a transition from a metastable pre-fusion state to a stable post-fusion state^26^. Theoretical studies have attempted to prevent RBD binding by inhibiting ACE2 receptor using different molecules^27–31^. Our previous studies have shown that plant-based molecules inhibit receptors and several proteins in the pathway by restricting conformational freedom^32^. Currently, symptomatic treatment is the only effective treatment option; therefore, the development of specific targeted drugs is crucial. Coronavirus ‘WH-Human 1’ proteomics sequences have recently become available as a result of metagenomic RNA sequencing of a patient in Wuhan. WH-Human 1 shares 89.1% of its DNA with a group of coronaviruses known as SARS coronaviruses^33^. Potential inhibitory antibodies can be identified by scanning millions of antibody sequences and detecting neutralizing antibodies based on this sequence. Antibodies neutralize SARS-CoV-2 by binding to either RBD or NTD, disrupting the interaction of the virus with human angiotensin-converting enzyme 2^34–37^. This mutation results in an allosteric conformational change in the RBD domain, thereby increasing the number of hydrogen bonds formed at the ACE2-RBD interface^38^. The antibody attaches to the RBD and competes with the ACE2 receptor. It can bind to multiple sites on the RBD, inducing conformational changes that hinder ACE2 binding, and it can also attach to the RBD, impeding the required conformational transition for effective viral protein binding^37^.

Previous studies have demonstrated that powerful antibodies interact with the ACE2-RBD and spike glycoproteins, resulting in decreased binding affinity or transmissibility^36–38^. The mechanisms by which conformational dynamics are propagated to other protein regions and how this knowledge can be used to design small-molecule drugs or therapeutic candidates remain largely unexplored. In this study, we utilized microsecond all-atom simulations to investigate the conformational changes induced by multiple RBD mutants when bound to ACE2 receptors and to elucidate how these changes are propagated to the receptor. Our study also demonstrated that the binding of the B38 monoclonal neutralizing antibody to the RBD domain restricted conformational fluctuations in the ACE2-RBD complex. Overall, our findings provide new insights into the ACE2-RBD interaction and the role of mutations in controlling receptor conformational flexibility of the ACE2 receptor.

### Computational Details

The crystal structure of the spike receptor-binding domain bound with angiotensin-converting enzyme 2 was obtained from PDB: 6M0J^39^. Glycan and ions from the crystal structures were maintained and the system was neutralized with 140 mM NaCl. Mutants were created using the 6M0J structure in the Chimera software package (Table 1). The dynamics of the wild structure (6M0J) were compared with a mutant simulation to elucidate the structural dynamics. The Charmm36 force field with the TIP3P water model^40, 41^ was employed for all simulations. Unbiased all-atom molecular dynamics simulations for 1 μs were conducted using GROMACS 2021v^42^. Visualization of the simulations was performed using the Chimera software along with matplotlib^43, 44^. Electrostatic interactions were computed using the particle mesh Ewald method^45^with a Verlet cutoff distance of 1.4 nm for short-range repulsive and attractive interactions. The temperature and pressure were regulated at 310 K and 1 bar by employing the Nose-Hoover temperature coupling and the Parrinello-Rahman barostat algorithm^46–47^, respectively. The LINCS algorithm^48^ was applied to constrain all bond lengths, and the simulation trajectories were saved every 100 ps with a time step of 2 fs. Before the production run, minimization with the steepest descent with a maximum of 5000 steps and equilibration was performed in both the NVT and NPT ensembles for 10 ns and 20 ns, respectively. All mutant systems were subjected to the same simulation protocol. Trajectory analysis was conducted using GROMACS tools and Python codes from the MDanalysis package^49^.

The Umbrella sampling method was employed to calculate the free energy associated with the unbinding of the RBD from the ACE2 receptor. Umbrella sampling is an efficient sampling technique that can be easily implemented with minimal modifications to existing MD simulation frameworks. To calculate the binding energy between ACE2 and RBD, the reaction coordinate considered was the separation between the center of mass of RBD and ACE2 along the x-axis. A simulation box was constructed to satisfy the minimum image convention, and the RBD domain was pulled 5 nm away from the center of mass of ACE2. The initial structures of both the apo and mutant ACE2-RBD complexes were minimized using the steepest descent method. Subsequently, 10 ns of NVT and NPT equilibration was performed, and the last frame from the NPT equilibration was used for the steered MD, employing a harmonic spring constant of 1000 kJ/mol and a pulling rate of 0.001 nm/ps. The RBD was gradually pulled to a distance of 5 nm and 500 frames were generated, each separated by a distance of 0.01 nm. Each frame was subjected to umbrella sampling simulation for 100 ns to obtain a smooth profile. The resulting one-dimensional PMF curve for each mutant provided ΔG, representing the binding energy, and the equilibrium probability distribution for all umbrella sampling windows and free energy profiles were obtained using the weighted histogram analysis method^50^ implemented in the GROMACS tool.

### Principal component analysis and free energy landscape

The independent collective motions of ACE2 and RBD were assessed independently using Principal Component Analysis (PCA) ^51^ to gain insights into their respective dynamics. To perform PCA, we constructed the variance-covariance matrix C based on the trajectories for an ensemble average:

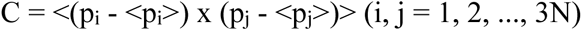

Here, p_i,j_ represents the mass-weighted coordinate vector of ACE2 or RBD comprising N atoms and <…> indicates the ensemble average. The coordinates of the backbone atoms were utilized to build the symmetric matrix C. We computed averaged values during the equilibrated phase of the simulations after aligning the trajectory to a reference structure using a least-squares fit procedure with the first frame to eliminate overall translations and rotations. The resulting matrix C is diagonalized by an orthogonal coordinate transformation matrix T, yielding a diagonal matrix R of eigenvalues λ_i_ as follows:

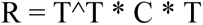

Each column of T represents an eigenvector describing the direction of motion relative to <p_i_>, and the eigenvalues in R represent the total mean square fluctuation of the system along the corresponding eigenvector. PCA was performed on the microsecond simulations using the MDAnalysis package in conjunction with the scikit-learn library^52^ to facilitate the efficient computation and comprehensive analysis of the principal components.

The free energy landscapes (FEL) of ACE2 and RBD were obtained using a conformational sampling method based on molecular dynamics simulation trajectories. This method allowed us to explore conformations near the native state structure^53, 54^. In our analysis, we focused on two important structural descriptors: the root mean square deviation (RMSD) and radius of gyrations along with the first two components of PCA. To achieve a three-dimensional representation of the FEL, we defined the probability of finding the system in a particular state characterized by a p_α_ value of RMSD and gyration as proportional to the exponential of the negative free energy:

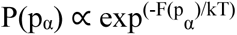

Where P(p_α_) is the probability of finding the system in a state with the specified RMSD and gyration values (p_α_). F(p_α_) represents the free energy of the state, k is the Boltzmann constant, and T is the temperature. A three and two-dimensional FEL was constructed by sampling multiple points along the RMSD and gyration reaction coordinates from the MD simulation trajectories. At each point, the corresponding free energy value was calculated using the negative logarithm of the probability distribution:

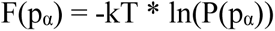

The resulting FEL provides valuable insights into the thermodynamics and kinetics of the conformational space explored by ACE2 and the RBD. This allowed us to visualize the distribution of states, identify stable conformations, and study the dynamics of mutants with the wild ACE2-RBD and apo ACE2 state structure.

### Clustering analysis

Simulation trajectories were employed to cluster the structures using the k-means clustering algorithm^56^, in combination with spectral clustering. The k-means clustering spectral clustering methods are widely used in unsupervised machine learning to segregate a set of data points. In this process, each frame is iteratively assigned to its closest centroid and the centroids are updated until convergence is achieved. Spectral clustering^57^ relies on the eigenvectors and eigenvalues of a similarity matrix constructed from the trajectories. The data were transformed into a lower-dimensional space by using the eigenvectors and eigenvalues of the Laplacian matrix of the similarity matrix. The final clustering was performed by applying k-means clustering to the transformed information using the first eigenvectors with the largest eigenvalues. Data were divided into five distinct clusters using the k-means clustering algorithm. To ensure robustness, the algorithm was executed ten times with different initial centroid seeds, and the maximum number of iterations for each run was set to 300. The combined approach of spectral k-means clustering led to improved clustering results. Spectral clustering played a crucial role in reducing the dimensionality of the trajectory and capturing its global structure, whereas k-means clustering further refined the analysis based on specific features, enhancing the clustering analysis. The simulation trajectory was processed using the MDAnalysis module and clustering analysis was performed using the scikit-learn library. This approach facilitated the identification and characterization of different conformational states within the ACE2 and RBD structures. By analyzing the cluster assignments and centers, valuable insights were obtained regarding the behavior and structural dynamics of ACE2 and RBD.

## Results and Discussion

To understand the structural transitions of the receptor-binding domain and mutation-induced transitions to the ACE2 receptor, microsecond all-atom molecular dynamics simulations were performed. The stability of the mutant conformation and movements of conformational transitions through the RBD were also evaluated. In addition, ACE2 conformational transitions from the apo, closed, and RBD-bound states were evaluated using 10 µs simulations. Additionally, we reported restricted conformational transitions of the RBD upon binding with B38-neutralizing antibodies.

### Conformational stability of Angiotensin-converting enzyme 2

Mutations in the RBD have been shown to exhibit a higher affinity towards the ACE2 receptor, which increases the transmissibility of the virus. To determine the underlying cause of this higher affinity, we conducted one-microsecond all-atom molecular dynamics simulations of both wild and mutants of RBDs with ACE2. Understanding the structural influence of these mutations towards the ACE2 receptor is crucial for characterizing binding patterns and conformational dynamics. The binding nature of ACE2-RBD using molecular dynamics simulations, and many have argued that strong interactions occur via hydrogen bonding and electrostatic interactions at the binding site^57–61^. Earlier studies indicate that the higher free energy contribution of the interacting residues is responsible for the residues located outside the receptor binding domain^62–63^. The root mean square deviations of backbone atoms and root mean square fluctuations of Cα of the ACE2 receptor upon binding to the wild-type and mutants are shown in Figures S1 and S3a. Higher RMSD and RMSF values were observed for the ACE2 receptor bound to mutants. The average backbone deviation of ACE2 from the wild RBD was found to be 0.23 nm whereas, mutants show a higher value of 0.6 nm for alpha- and kappa mutants. Higher fluctuations in ACE2 receptors indicate that structural transformation or steric hindrance induces conformational transition. These conformational changes induced by the binding of the mutant RBD were further investigated using principal component analysis. The PCA-based backbone atoms are denoted by the eigenvectors of the covariance matrix, which are indicated by their coincident eigenvalue and the total concerted motion of the protein, and are correlated with protein functions. Figure S4a shows the motion along the first and second components of ACE2 backbone atoms. The alpha variants showed higher fluctuations in the region between 600-1700 atoms, corresponding to the α-helix chain from Ser19-Asn103, which is involved in RBD binding. The Ser19-52Thr and 55Thr-82Met regions are primary α-helix bundles that interact with the RBD and N-terminal domain of the ACE2 receptor. These fluctuations can be categorized as primary fluctuations because they arise from direct interactions with the RBD. The beta and epsilon mutants showed higher fluctuations in the domain Gln89-Gln102, which is a small helix bundle situated behind the binding site. These fluctuations are characterized as secondary fluctuations because they arise as a consequence of the primary fluctuations in the helix bundles of Ser19-52Thr and 55Thr-82Met. The time evolution of the secondary structural changes of ACE2 receptors was obtained from a one-microsecond simulation and is depicted in Figure S5. The initial conformation of ACE2 receptors showed stable dynamics and a transition from 63.3% helix, 3.7% sheet, 8.4% turn in the first frame, and 24.7% coil to 60.7% helix, 3.5% sheet, 6.2% turn, 28.9% coil, and 0.7% 3-10 helix at the last frame of 1 µs simulation. The alpha variants show the occasional transition of helix, coil, and bend between residues 200-300 and the final secondary content shows 57.1% helix, 3.6% sheet, 10.9% turn, 27.6% coil, and 0.9% 3-10 helix. These residues are located within the N-terminal region of the RBD, which lies outside the binding domain and exhibits fewer interactions with ACE2. Secondary and tertiary structural transitions occurred in these regions, resulting in higher fluctuations in ACE2. The secondary structural changes in ACE2 receptors in the wild and mutant systems are shown in Table S17. Transitions from the α-helix to occasional bends and turns, along with the formation of small β-sheets, were observed in all the mutants. In addition, tertiary structural changes in the receptor were evaluated by constructing a minimum-distance matrix (Figure S6). The kappa mutant showed the highest deviations in both RMSD matrices, but there were no significant variations in the distance matrix, indicating possible tertiary structural stability of ACE2 upon binding to the RBD. This further indicates that the RMSD fluctuations of the backbone and PCA-based fluctuations of ACE2 receptors arise from minor secondary structural changes, as well as fluctuations at the binding site. The formation of hydrogen bonds and salt bridges at the binding site of ACE2-RBD also leads to further structural fluctuations in the RMSD and PCA.

### Conformational dynamics of receptor binding domain (RBD)

Understanding the conformational dynamics of the receptor-binding domain is crucial for deciphering the ACE2 binding mechanism. The structural stability was characterized by assessing the RMSD, Cα RMSF, and PCA-based fluctuations of the backbone atoms. Because mutations occur in the RBD, higher structural fluctuations and conformational changes were prominent than ACE2 (Figure S2). RMSF of the Cα atoms showed a substantial increase in the mutant RBD, as depicted in Figure S3b. The highest fluctuations were observed between residues 104-157, which were located at the binding site. All mutations showed the rearrangement of residue side chains at the binding site and the formation of hydrogen bonds with ACE2 receptors. The formation of a higher hydrogen-bonding network at the ACE2-binding sites is crucial for a strong affinity^42^. The PCA-based backbone fluctuations of the RBD were obtained to understand the specific fluctuations at the binding site (Figure S4b). The alpha mutations that represent N501Y showed a wide peak in the PCA backbone corresponding to the residues Glu354-Lys357. Interestingly, these residues were far from the binding site, as well as from N501Y. This suggests that point mutations at position 501 triggered conformational changes in the RBD, which propagated to other regions. The secondary structure showed 7.7% helices, 28.9% sheets, 17.5% turns, and 45.9% coils at the end of the simulation, corroborating this transformation. The starting conformation had 6.2% helix, 30.9% sheet, 17.5% turn, 42.3% coil, and 3.1% 3-10 helix, and the formation of minor helical structures further enhanced the conformational stability of the RBD (Figure S7 and S8). The Free-energy landscape (FEL) obtained from the first two principal components of PCA revealed increased structural stability in the presence of N501 mutations (Figure 1). FEL showed broad and deep minima for the wild-type RBD, whereas the N501 mutation indicated the presence of multiple evenly distributed minima, indicating various conformational changes in the RBD. This suggests that there were fewer conformational transitions in the RBD upon ACE2 binding. The mutation E484K, along with N501Y, induces more conformational transitions and high fluctuations in the Phe373-Phe377 and Asp389-Cys391 regions, which are opposite ends of the binding site. We observed β-sheet formation along with occasional meta-states of coil-bend transitions in these regions. The free-energy landscape of the double mutant revealed the presence of several distinct wells with three minima associated with the lowest fluctuations. Notably, the transition from a single-point mutation to multiple mutations (beta to omicron) resulted in significant variations in the free-energy landscape, with multiple spikes in the PCA to a single wide spike observed in the omicron mutant. The region Leu387-Leu390 shows the highest fluctuations for K417N, E484K, and N501Y; these triple mutants showed multiple transitions from α-helices to coils through random coils and bend regions. Higher transitions from the N-terminal region to the binding site indicate secondary structural changes in the mid-region of the RBD. The gamma variants with K417T, E484K, and N501Y mutations showed structural changes similar to those of the RBD. However, 417T caused a significant loss in the α-helix and the formation of bends and coils. The PCA-based backbone fluctuations were residue-specific, whereas beta variants exhibited a wide range of fluctuations. The beta variant showed 6.2% helix, 26.8% sheet, 22.2% turn, and 44.8% coil, whereas the gamma variant showed 3.1% helix, 28.4% sheet, 20.6% turn, 45.9% coil, and 2.1% 3-10 helix. The transformation of α-helices to β-sheets or loss of helical content changed the hydrogen-bonding pattern between the residues in the RBD domains, leading to higher fluctuations, as observed in the RMSD and RMSF values. Furthermore, the free-energy landscape of the beta mutant showed two wide minima, whereas that of the alpha variants showed multiple minima with sharp peaks. The 417T mutation further stabilized the RBD domain through structural changes, as observed in the free energy landscape. This indicates a well-connected wide minimum at the centre of the FEL, as depicted in Figure 1. Kim et al reported the differential interaction of ACE2-RBD interactions and mutants of concern. Their study of alpha, beta, gamma, and delta variants showed a higher pulling for the detachment of the RBD from ACE2 receptors due to N501Y mutations, in addition to the role of N90-glycan, followed by beta, gamma, and delta variants. They also reported a higher number of contact residues in RBD mutants with ACE2 receptors. Here, we observed fluctuations in certain residues far from the mutation sites, which caused secondary structural changes in different parts of the receptor. The RBD domain, which interacts with the S2 domain, showed significant variations in alpha-helical content. It travels through proteins and imitates the tight binding to the ACE2 receptor. The delta variant with L452R and T478K mutations at the binding site causes a maximum of 0.15 nm fluctuations, which is 70 % less than that of other variants. In the delta variant, 6.7% helix, 33.0% sheet, 19.1% turn, 38.1% coil, and 3.1% 3-10 helix content were observed. Consequently, the free-energy landscape showed a single wild well with a single minimum that diverged from a wider well. Here, a transformation from coil to bend and turn, along with occasional α-helices and β-sheets, is observed in the secondary structure. In the case of multiple mutations compared to a single mutation of L452R (epsilon), the secondary structure content shows a 10.3% helix, 27.3% sheet, 17.5% turn, 42.8% coil, and 2.1% 3-10 helix. A greater loss of secondary structure was observed in the C-terminal region of RBD, with multiple minima spread across the principal components, as shown in Figure 1q. The double mutations L452R and E484Q (kappa) could potentially have higher transmissibility, severity, or reinfection risk and require continuous monitoring^64, 65^. Kappa mutations are highly contagious and can evade neutralizing antibodies^64^. The free-energy landscape of the kappa variant exhibited a wide peak with the lowest energy and a single spike, as shown in Figure 1i. The variant showed a 6.2% helix, 29.4% sheet, 13.9% turn, 47.9% coil, and 2.6% 3-10 helix in its secondary structure and formed a α-helix and turn region at the C-terminal region. The lambda variant with L452Q and F490S mutations also showed a similar two-peak FEL, but it was found that both peaks were moderately wide and multiple peaks were observed in the backbone atoms, especially in the region Tyr380-Phe400. The α-helix in the N-terminal region, which is situated at the binding site of ACE2, transforms to bend and turn. Multiple meta-states were also observed between the turn-coil bend, which resulted in 6.7% helix, 30.9% sheet, 25.8% turn, and 36.6% coil in its secondary structure. The omicron mutation, which reported the highest number of mutations (G339D, S371L, S373P, S375F, K417N, N440K, G446S, S477N, T478K, E484A, Q493R, G496S, Q498R, N501Y, and Y505H), showed fluctuations in residues between Asn450-Thr500. The free-energy landscape of the omicron variant exhibited a wide minimum among all the mutants. The secondary structure transitioned to 9.3% helix, 32.6% sheet, 22.3% turn, and 35.8% coil structures. These mutations result in higher structural stability of the RBD domain. This is possible through conformational transitions and inter-residue interactions with ACE2 receptors. The mutations not only lead to higher affinity and transmissibility but also transform the RBD into more energetically stable conformations, as observed in the free energy landscape. Interestingly, these mutations cause structural changes in the RBD that are distant from the binding and mutation sites.

**Figure 1.**
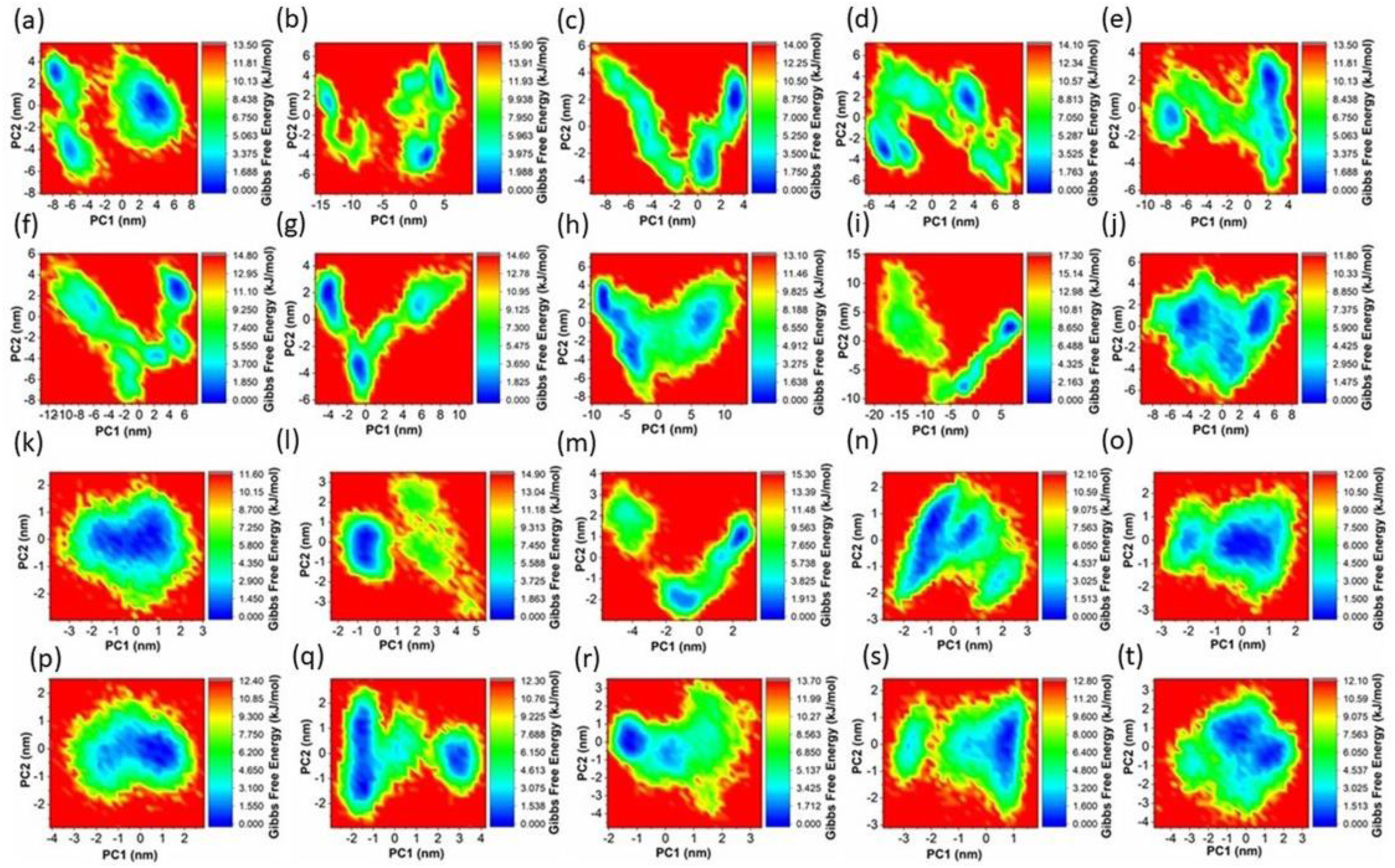
2D-free energy landscape (FEL) was constructed from the first and second principal components. a-j represents the ACE2 receptor, and k-t represents the RBD. (a)wild,(b)alpha,(c)alpha+,(d)beta,(e)gamma,(f)delta,(g)epsilon,(h)kappa,(i)lambda, and (j)omicron. (k)wild,(l)alpha,(m)alpha+,(n)beta,(o)gamma,(p)delta,(q)epsilon,(r)kappa,(s)lambda, and (t)omicron. The wild ACE2 receptor shows deep and wide global minima, along with well-connected local minima. However, interaction with the mutant RBD led to several deep local minima, and the omicron mutant showed global minima with multiple wells. This indicated the strong affinity of the RBD omicron variant for the ACE2 receptor. Interaction with this variant causes the ACE2 receptor to reach a global minimum. In RBD, the wild type shows global wide minima, and among the variants, the stability increases when the number of mutations increases, and the maximum is identified in the case of the omicron mutant.

### ACE2-RBD interactions

The binding affinity between angiotensin-converting enzyme 2 (ACE2) receptor and receptor-binding domain (RBD) is primarily driven by multiple intermolecular forces, including hydrogen bonding, electrostatic interactions, van der Waals forces, and salt bridges^66–69^. The hydrogen-bonding network at the binding interface between wild-type ACE2 and RBD consisted of an average of nine hydrogen bonds, which contributed significantly to the stability of the complex (Figure S9a). However, upon analysis of the mutations, excluding the omicron variant, the hydrogen-bonding network was reduced to an average of seven hydrogen bonds. In contrast, the omicron variant exhibited the highest (9-10) average number of hydrogen bonds. Initial crystal structure analysis of the ACE2-RBD complex revealed 10 hydrogen bonds between Ser19-Ala475, Gln24-Asn487, Lys31-Gln493, Glu35-Gln493, Glu37-Tyr505, Asp38-Tyr449, Tyr41-Thr500, Gln42-Gln498, Tyr83-Asn487, and Lys353-Gly502. These hydrogen bonds were found to have a maximum energy of 25 kJ/mol and a total hydrogen bond energy of 196 kJ/mol. However, at the end of the simulation, residues Gln24-Asn487, Glu35-Gln493, Tyr-83-Asn487, Lys353-Gly502, and Asp355-Thr500 formed hydrogen bonds, with a total energy of 96 kJ/mol. The alpha mutation showed nine hydrogen bonds in the initial structure, similar to the wild type, but formed two hydrogen bonds with Lys353-Tyr501 and Lys353-Gln502. The final structure of the alpha mutation showed three hydrogen bonds with a total energy of 51 kJ/mol: Ala475-Gln24, Asn487-Tyr83, and Gly502-Lys353. An additional mutation in the alpha+ increases the total hydrogen bond strength to 185.08 kJ/mol with interactions such as Ser19-Ala475, Gln24-Asn487, Lys31-Gln493, Glu35-Gln493, Glu37-Tyr505, Tyr41-Thr500, Gln42-Gln498, Tyr83-Asn487, and Lys353-Tyr501. However, these hydrogen bonds were restricted to five residues (Ser19-Ala475, Glu35-Gln493, Tyr41-Thr500, Tyr83-Asn487, and Lys353-Gly502), with a total energy of 108 kJ/mol. The addition of the K417N mutation did not increase the hydrogen bond energy (149 kJ/mol). However, the initial structure contained nine hydrogen bonds: Ser19-Ala475, Glu37-tyr505, Asp38-Tyr449, Tyr41-Th500, Glnn42-Gln498, Tyr83-Asn487, Lys353-Tyr501, and Lys353-Gly502. These hydrogen bonds decreased at the end of the simulations (Ser19-Ala475, Glu37-tyr505, Asp38-Tyr449, Tyr41-Thr500, Gln42-Gln498, Tyr83-Asn487, Lys353-Tyr501, and Lys353-Gly502). The delta mutations showed higher hydrogen bond energies for both the initial and final frames, the residues Ser19-Ala475, Lys31-gln493, Glu35-Gln493, Glu37-Tyr505, Asp38-Tyr449, Tyr41-Thr500, Gln42-Gl498, Tyrr83-Asn487, and Lys353-Gly502; the final structure showed a hydrogen bond between the residues Ser19-Ala475, Gln24-Ala475, Lys31-Glu484, Tyr83-Asn487, Glu329-Arg439, Lys353-Gly502, and Asp355-Thr500. When it moved from the alpha to the omicron, the residues that participated in the hydrogen bond patterns were similar when they moved from the alpha to omicron. The mutation induces structural rearrangement, which reduces the distance between the hydrogen bond acceptor donors and increases the energy between these residues. The omicron mutations exhibited the lowest average distances between residues to form hydrogen bonds with Ser19-Ala475, Gln24-Asn487, Asp38-Ser496, Asp38-tyr449, Tyr41-Thr500, Gln42-Arg498, Tyr83-Asn487, Lys353-Ser496, and Lys353-Gly502, with the highest energy of 230 kJ/mol (Figure S10 and S11). Interestingly, the native ACE2-RBD showed an average electrostatic interaction energy of −285 kJ/mol, whereas the omicron variants show −480 kJ/mol (Figure S10b). These significant differences in the electrostatic energy at the binding site arise from changes in the distance between the residues arising from structural rearrangements. The initial changes observed in the mutant static structure were consistent throughout the simulations. The collective movements of the Cα atoms of both RBD and ACE2 indicate conformational waves through the protein due to mutations at different sites. The movements in the RBD arise due to mutations, whereas ACE2 fluctuations arise due to the induced conformational changes resulting from the strong binding of the RBD. However, the average van der Waals interaction energies between ACE2-RBD were found to be lower for the mutants than for the wild type. The binding sites of ACE2 consist of charged, uncharged, and polar amino acids such as Ser19, Gln24, Lys31, Glu35, Glu37, Asp38, Tyr41, Gln42, Tyr83, Lys353, Ala475, Asn487, Gln493, Tyr505, Tyr449, Thr500, Gln498, Asn487, and Gly502. The solvent-accessible surface area increased from alpha to omicron mutants, as shown in Figures S12-15. Furthermore, we calculated the salt bridge between ACE2-RBD using a 4 Å cut-off distance. The native ACE2-RBD contains 112 salt bridges over 10000 frames. In contrast, the mutants showed more salt bridges, and omicrons showed the highest number of salt bridges among the mutants (197 salt bridges). This indicates that salt bridge formation between ACE2-RBD also plays a crucial role in the stabilization of binding. To gain further insight into the dissociation events between ACE2-RBD, we performed umbrella sampling simulations to calculate the potential of the mean force (PMF) for complete dissociation of RBD from ACE2 (Figure S16). The wild-type complex showed a dissociation energy of 17.82 kcal/mol, whereas the omicron variant showed the highest dissociation binding energy of 25.10 kcal/mol. Variations in energy arise owing to the differences in the interacting residues. ‘Conformational waves’ travelling from the mutant site to the binding site induced conformational changes, and rearrangement of the residues at the binding site led to different dissociation energies, which led to higher residue interactions. It is crucial to understand ACE2 dynamics at different states to decipher the RBD-induced binding pattern through long-time simulations. Therefore, we evaluated ACE2 dynamics in the native open and closed states using 10 µs all-atom simulations.

### Dynamics of Angiotensin-converting enzyme 2 and B38 neutralizing antibody

The dynamics of ACE2 receptor were evaluated using 10 µs all-atom simulation trajectories^70^ of five structures: apo open state^71^ (PDB ID:1R42), inhibitor-bound closed state^71^ (PDB ID:1R4L), complex state with the receptor-binding domain of a spike protein from SARS-CoV-1 (PDB ID:2AJF)^72^, complex state with the receptor-binding domain of a chimeric construct of SARS-CoV-2 (PDB ID:6VW1)^73^, and complex state with the receptor-binding domain of a spike protein from SARS-CoV-2 (PDB ID:6M17)^74^. Here, we mainly emphasized PCA and FEL to comprehend the conformational changes occurring in the ACE2 receptor throughout the 10 µs simulation (Figure 2). These motions allowed us to identify the principal structural features responsible for the changes induced by RBD, and to determine the highly flexible or rigid regions of the protein that contribute to these features. PCA can also be used to identify collective motions that occur in a protein, such as the opening or closing of a protein domain or the movement of a protein along a particular axis. By identifying these collective motions, a better understanding of the functional characteristics of the receptor and its interaction with RBD can be achieved. The unbound state of the receptor exhibited a collective representation of the principal components in a 2D projection. Moreover, 3D projection of the free-energy landscape revealed stable receptor dynamics with global minima. In contrast, the closed conformation displayed increased fluctuations in the presence of the local minima, eventually resulting in stable positions. PCA components revealed the distribution of eigenvectors across a wide range relative to the apo state. CoV1 and CoV2 (6VW1) exhibit multiple minimum and high receptor fluctuations, respectively. Conversely, the 6M17 structure exhibited a global minimum, with fewer conformational transitions. Binding of RBD stabilizes the receptor earlier than the apo state, as shown by the free energy landscape. Furthermore, the distribution of RMS fluctuations revealed a stable distribution of backbone fluctuations, as illustrated by the violin plot of RMSD. The binding of the COV2 receptor results in the stabilization of the ACE2 receptor and possible fast signal transduction, which may lead to faster transmissibility. Additionally, clustering analysis of the ACE2 receptor revealed that the unbound protein exhibited a larger number of clusters (520) than the closed, CoV1, 6VW1, and 6M17 forms, which exhibited 192, 122, 126, and 56 independent and diverse conformations, respectively (Figure 3). These results suggest that RBD binding restricts the conformational freedom of the receptor. Cluster analysis of the ACE2 apo form revealed significant structural heterogeneity with 530 distinct structural clusters. In contrast, the ACE2-RBD complex contains only 56 clusters, indicating a much greater degree of structural homogeneity. The higher number of clusters observed in the ACE2 apo form suggested that the protein was structurally more flexible in its unbound state. This flexibility may be important for the protein to undergo conformational changes upon binding to the RBD and accommodate the RBD-binding interface. Furthermore, the structural heterogeneity observed in the ACE2 apo form may play a role in the promiscuity of viral entry as it may allow the protein to interact with various related and unrelated ligands. However, the lower number of clusters observed in the ACE2-RBD complex suggests that the interaction was more structurally rigid and less flexible. This rigidity may be important for stabilizing the complex and ensuring proper orientation of the RBD-binding interface with ACE2. The greater structural homogeneity observed in the ACE2-RBD complex may also explain why neutralizing antibodies and vaccines are primarily directed towards the RBD-binding interface. The human-origin monoclonal antibodies binding to the spike glycoprotein receptor binding domain of the virus can reduce virus titers in infected lungs and lower infectivity in vivo mouse models^75^. Our findings suggest that the B38 antibody-bound RBD undergoes fewer structural transitions than the apo state, likely because of the strong binding of the antibody, which restricts the conformational dynamics of the RBD domain (Figure 4). This strong binding results in decreased affinity for the ACE2 receptor, leading to reduced virus transmissibility, as observed in mouse models^75^. Here the apo state displays extensive residue movements, while the antibody-bound RBD demonstrates constrained movements. The free energy landscape of the antibody-bound RBD also suggests that the domain is at a deep global minimum during microsecond simulations, indicating the restricted movement of RBD residues (Figure 5). The clustering of backbone atoms further confirms that the apo state has a higher number of clusters than the antibody-bound state, indicating restricted movement of RBD residues. Interestingly, the RBD interaction with ACE2 showed slower fluctuations compared to antibody-bound RBD binding. More clusters were observed when RBD-nAbs interacted, suggesting that binding of neutralizing antibodies restricted RBD fluctuations, reduced transmissibility, and increased receptor movement, leading to higher structural flexibility (Figure S17 and S18). The free energy landscape with multiple local minima also indicated unstable receptor-RBD interactions due to the strong binding of the B38-neutralizing antibody.

**Figure 2.**
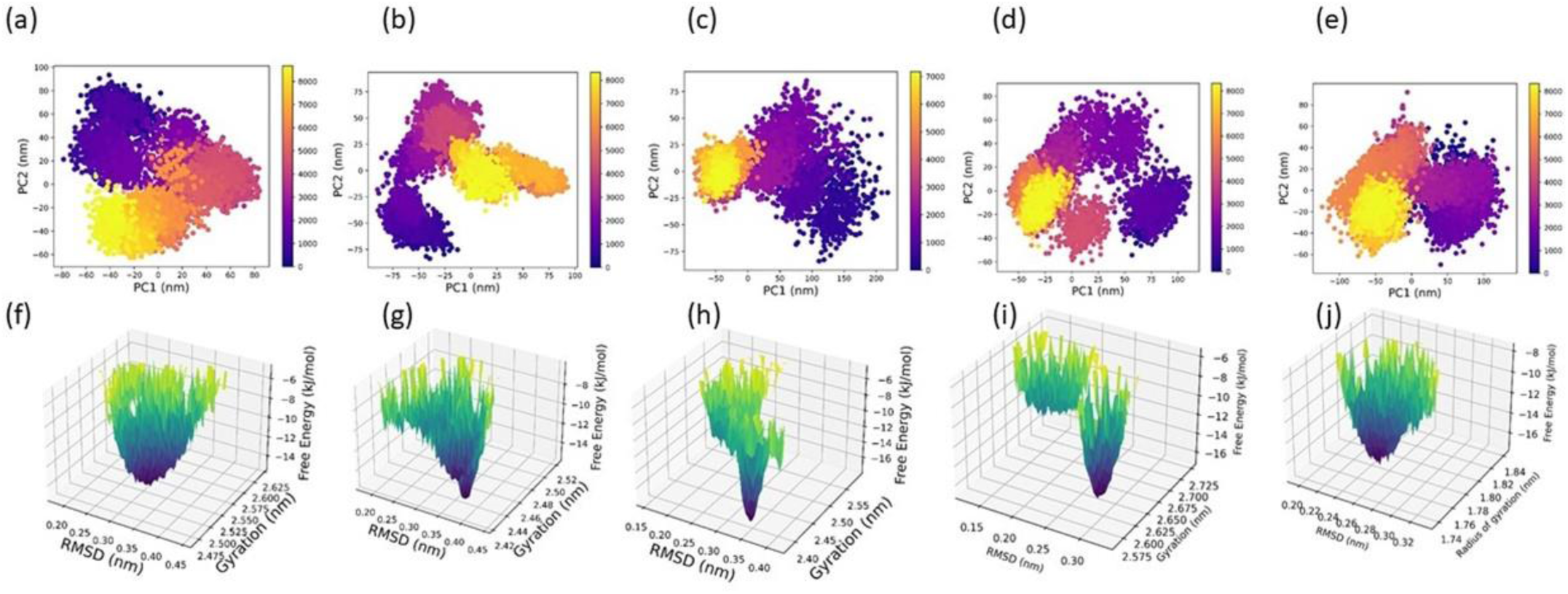
Distribution of principal components of the ACE2 receptor and free energy landscape. (a) apo, (b)closed,(c)cov1, (d)6vw1, and (e)6m17 represents 2d representation of the principal components. (f) apo, (g) closed,(h) cov1, (i) 6vw1, (j) 6m17 represents the 3-D free energy landscape obtained from the RMSD of backbone atoms and radius of gyrations. The apo state had an ordered and centered distribution of clusters in the PCA plot, whereas the closed state was centered and showed an uneven distribution. The cov1 showed an expanded distribution of clusters that extended beyond the ranges observed in the apo and closed structures. The cov2 structure possesses evenly distributed clusters but isolated and tightly clustered regions. The apo structure distribution of the points is ordered and oriented towards the center, which also suggests a high degree of correlation between the different principal components. The differences in the clusters also indicate that the conformations may interact with different ligands or molecules in different ways. The closed state indicates that the receptor underwent significant conformational changes, resulting in reduced stability. Binding of the RBD domains leads to higher flexibility and structural dynamics of the receptor. The apo state shows a wide distribution of FEL with a single well, indicating a conformational space and relatively low stability in terms of ligand binding and receptor action. The closed state showed a wide distribution, but sharp and well-represented a significant transition to a stable conformation. As expected, the ligand-bound receptor was more stable than the unbound apo receptor. The cov1 bound structure shows a wide distribution, but some parts have gone deep well, indicating that the receptor underwent a significant conformational change. The cov2 bound state (6vw1) showed multiple stable conformations with different levels of stability, whereas 6m17 showed a deep well with the possible receptor conformational space.

**Figure 3.**
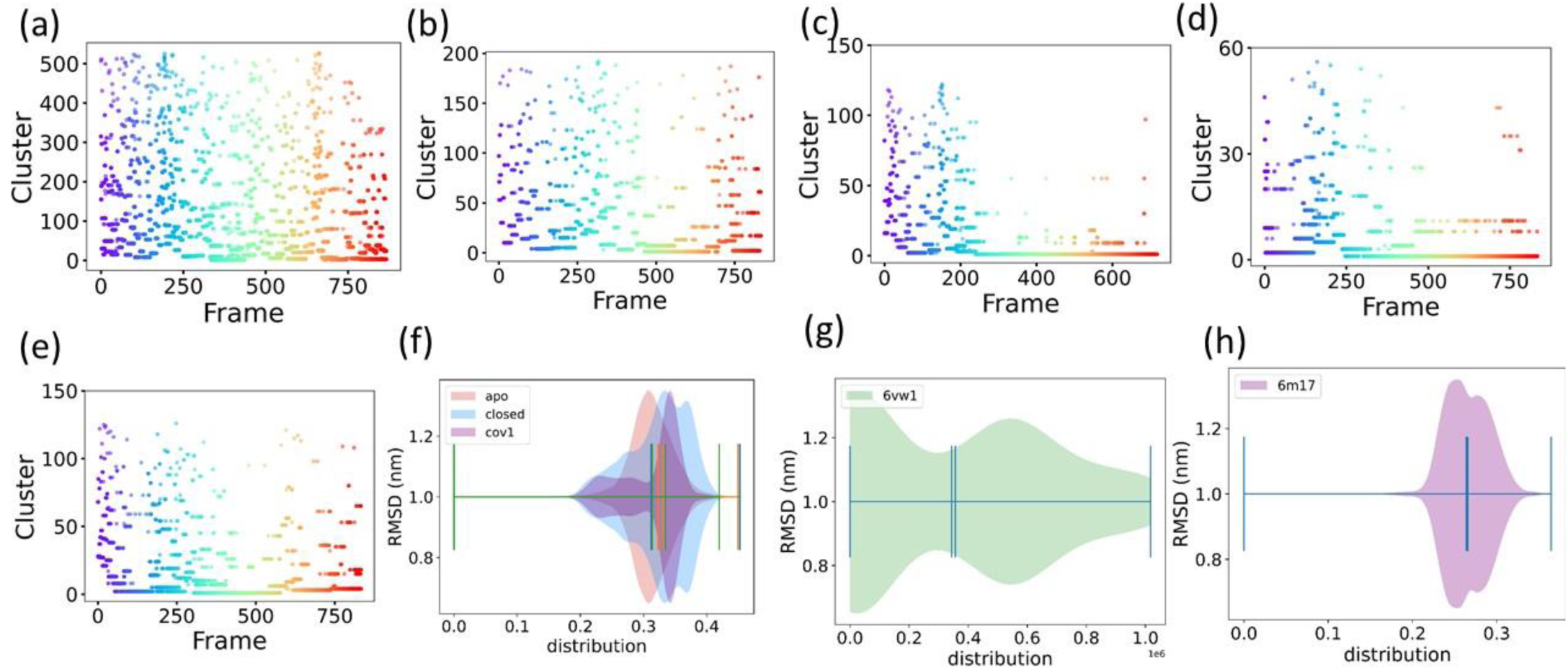
The clustering of ACE2 receptor (a) apo, (b) closed, (c) cov1, (d) 6vw1, (e)6m17 with corresponding backbone RMSD distribution. A higher number of clusters in the ACE2 apo structure indicates conformational diversity and structural states. This indicated that the unbound apo state of the receptor had a greater number of conformational states. Binding of the inhibitor and RBD domains reduced the conformational spaces due to strong binding. The higher conformational flexibility of the receptor increases the likelihood of interaction and leads to potential binding to different viral strains or mutants, which improves transmissibility. (f) - (h) represents the violin plot of RMSD distribution of corresponding ACE2 structures.

**Figure 4.**
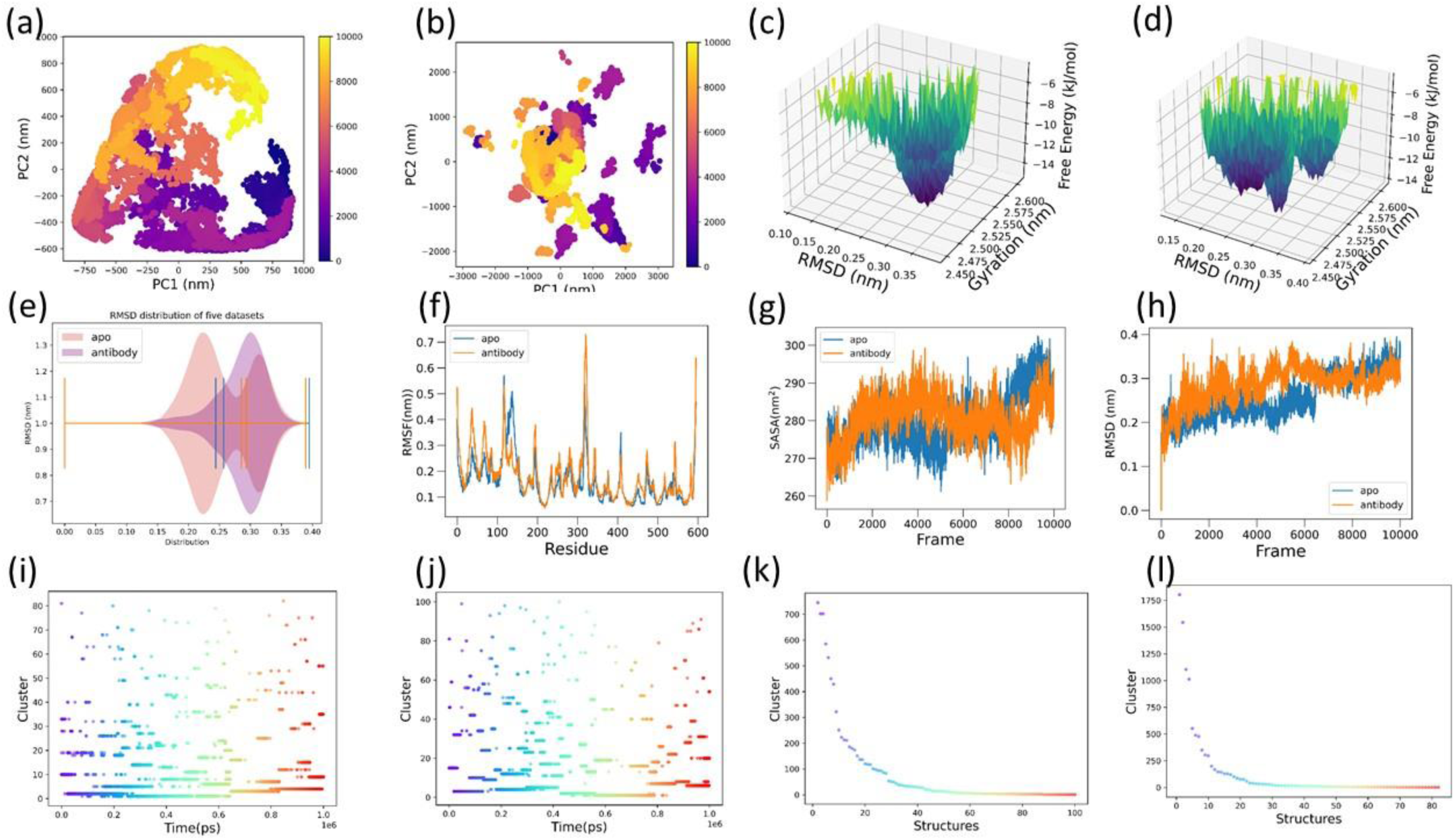
The dynamics of the ACE2 receptor bound to the antibody-inhibited RBD. (a) and (b) show the PCs distributions of the ACE2 receptor apo and RBD bound, respectively. The corresponding FEL generated from the RMSD and radius of gyration are given in (c) and (d). Changes in the RMS deviations and fluctuations are shown in (e), (f), and (h). (i)–(l) represent the clusters in each time frame and cluster per structure, respectively. The wide distribution of PCs components indicates a broader range of conformational states in the apo state, whereas ACE2, which binds to the RBD and is inhibited by the B38 antibody, suggests new conformational states and restricted certain motions that exist in the apo state of ACE2. This suggested that the antibody and its impact on the conformational dynamics of the ACE2-RBD complex. Antibody binding stabilizes or changes the binding regions of the RBD and reduces the normal binding pattern with the ACE2 receptor. Furthermore, the FEL suggested a stable and wide ACE2 conformation, which was restricted by the RBD by introducing multiple local minima states. In addition, the binding of RBD was found to alter the clusters and transformations of backbone atoms.

**Figure 5.**
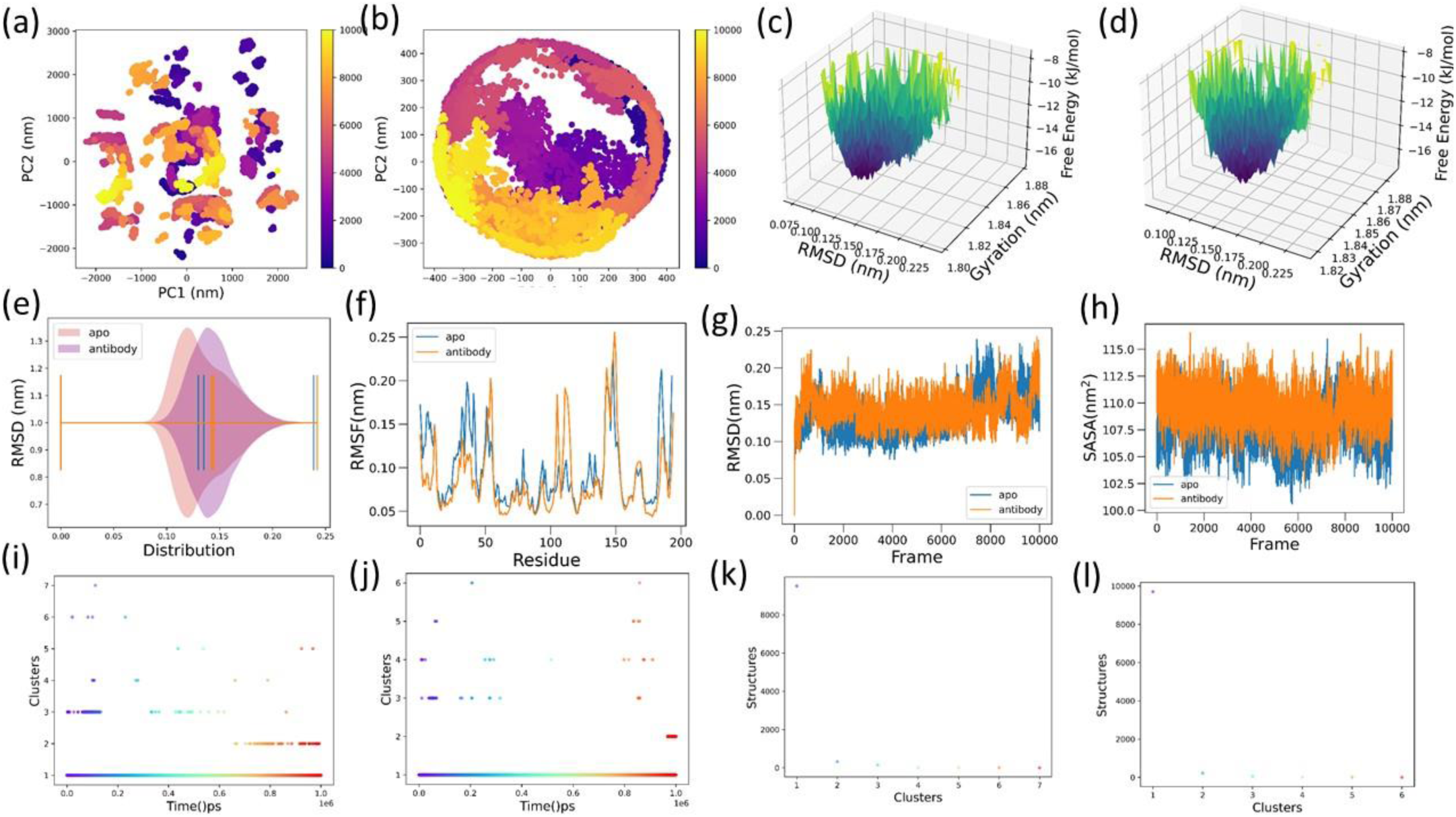
RBD dynamics with neutralizing antibodies and the ACE2 receptor. (a) PC representation of RBD bound to neutralizing antibody. (b) PCs representation of the RBD with ACE2 receptor, which is inhibited by neutralizing antibodies. (c) and (d) represent the corresponding free energy landscapes obtained from the RMSD and radius of gyration, respectively. RMS distribution (e), RMSF (f) RMSD (g), and solvent-accessible surface area (h) of RBD bound to the antibody. (i)–(l) represent the cluster per time and cluster per RBD structure.

## Conclusion

We present the dynamics of ACE2-RBD using long-time all-atom molecular dynamics simulations. The structural stability and dynamics of the RBD in various ACE2 receptor variants (wild-type, alpha, alpha+, beta, gamma, delta, lambda, kappa, epsilon, and omicron) induce conformational transitions from the RBD to the ACE2 receptor. Conformational changes arise from a single point mutation or multiple mutations in the RBD that determine the binding affinity. Conformational waves from the point mutation site travelled through the RBD to reach the ACE2 receptor. These conformational waves changed the orientation of the RBD residue at the binding site. Furthermore, the dynamics of ACE2 in its apo, open, closed, and RBD-bound states revealed induced transformations and the structural stability of the ACE2 receptor; RBD binding restricts the conformational freedom of the receptors, which prevents the binding of the designated ligands.ACE2 receptor shows significant structural heterogeneity, whereas it’s binding to the RBD domain indicates a much greater degree of structural homogeneity; the receptor exhibits greater flexibility in its unbound state, but binding of RBD domains induces structural transitions; the structural heterogeneity observed in the ACE2 unbound form plays a role in the promiscuity of viral entry, as it may allow the protein to interact with various related and unrelated ligands, and rigidity may be important for stabilizing the complex and ensuring the proper orientation of the RBD-binding interface with ACE2. The greater structural homogeneity observed in the ACE2-RBD complex revealed the effectiveness of neutralizing antibodies and vaccines that are primarily directed towards the RBD-binding interface. The binding of the B38 antibody revealed restricted conformational transitions in the RBD and ACE2 receptors owing to the tight binding of the monoclonal neutralizing antibody.

## Associated Content

### Supporting Information

Additional figures for ACE2-RBD wild, mutants and PMF.

## Acknowledgement

The author thanks RIKEN R-CCS Supercomputer Fugaku for the simulations through project hp210295.

---

**Figure S1.**
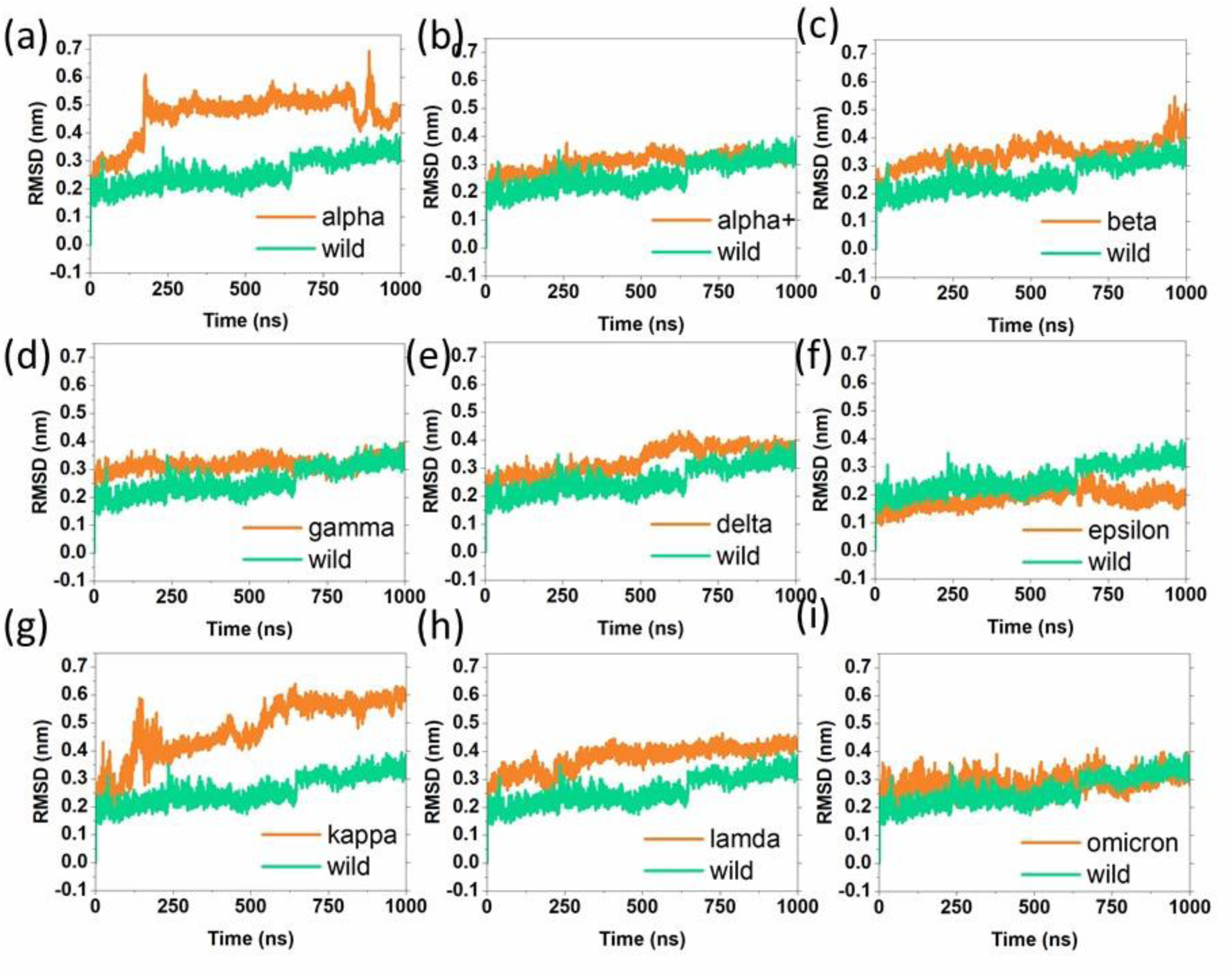
The root mean square deviations (RMSD) of ACE2 receptor backbone atoms when it bound with RBD mutant (a) alpha,(b)alpha+,(c)beta,(d)gamma,(e)delta,(f)epsilon,(g)kappa,(h)lambda, and (i) omicron structures respectively. The induced structural transitions from RBD to ACE2 resulted in higher RMSD

**Figure S2.**
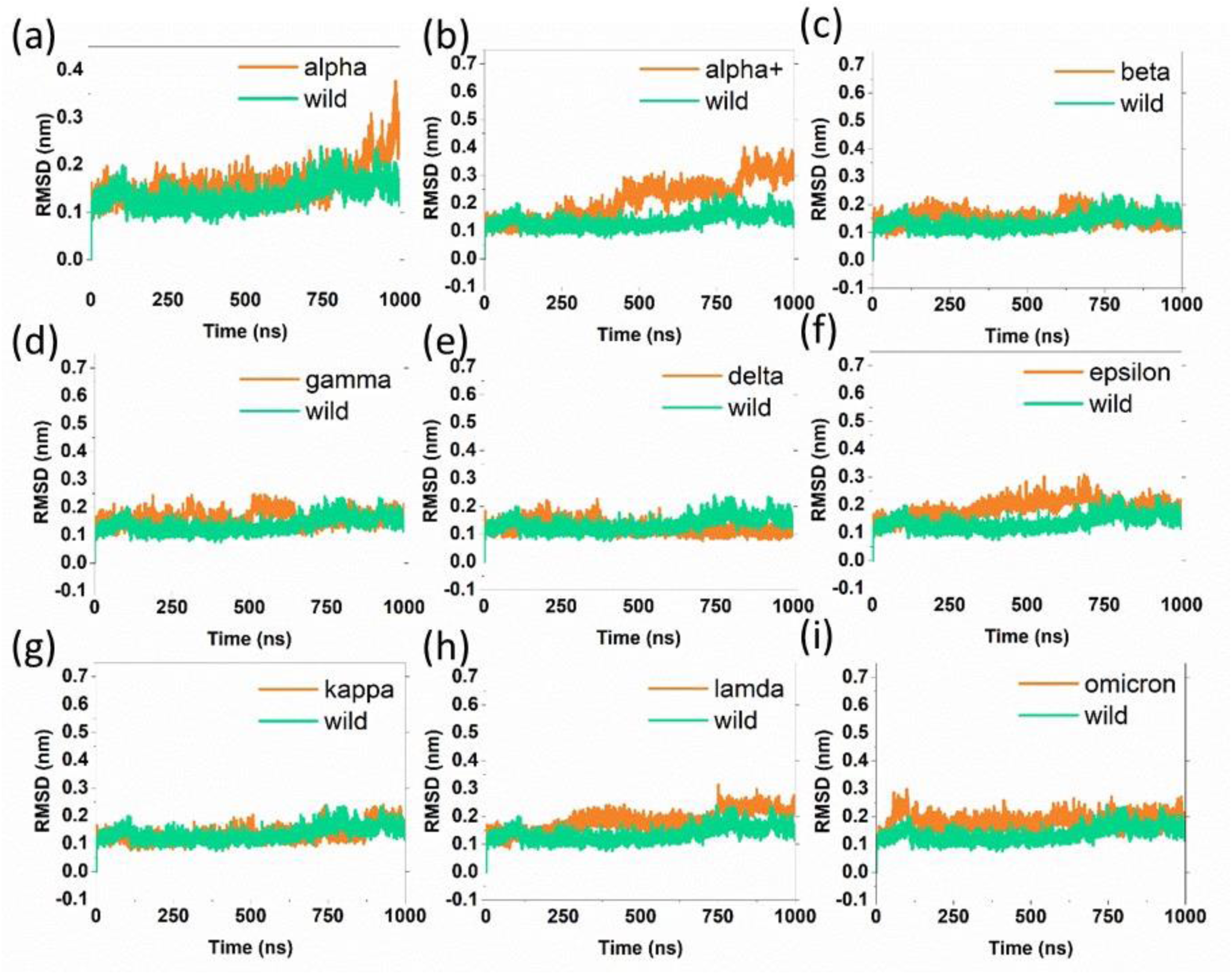
The root mean square deviations (RMSD) of RBD backbone atoms when they bound with ACE2 receptor (a) alpha,(b)alpha+,(c)beta,(d)gamma,(e)delta,(f)epsilon,(g)kappa,(h)lambda, and (i) omicron. The mutations induce higher RMSD fluctuations in mutants compared to the wild.

**Figure S3.**
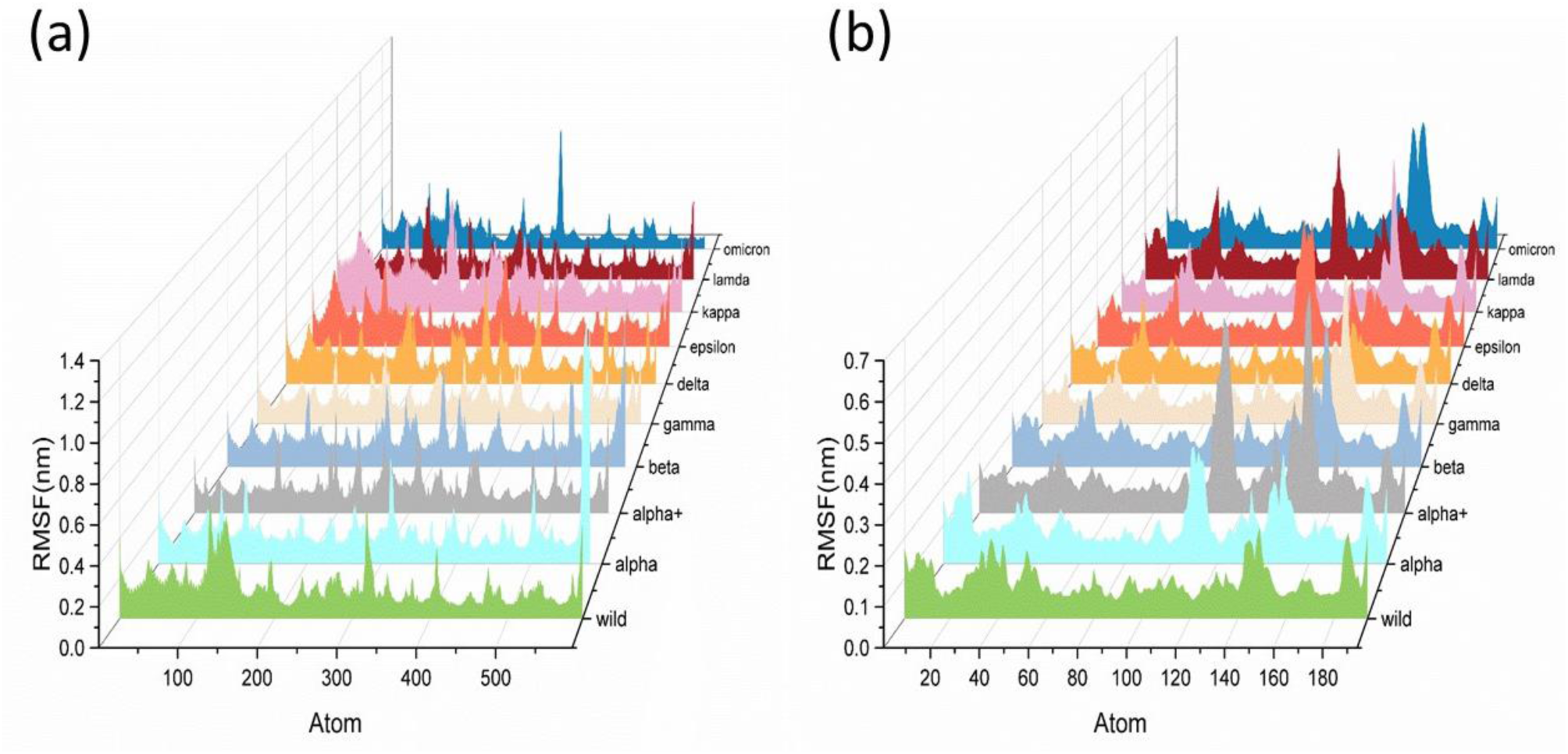
The root mean square fluctuations (RMSF) of Cα atoms of (a) ACE2 receptor and (b) RBD wild and mutants. The ACE2 receptor and RBD mutant show higher RMSF compare to the wild.

**Figure S4.**
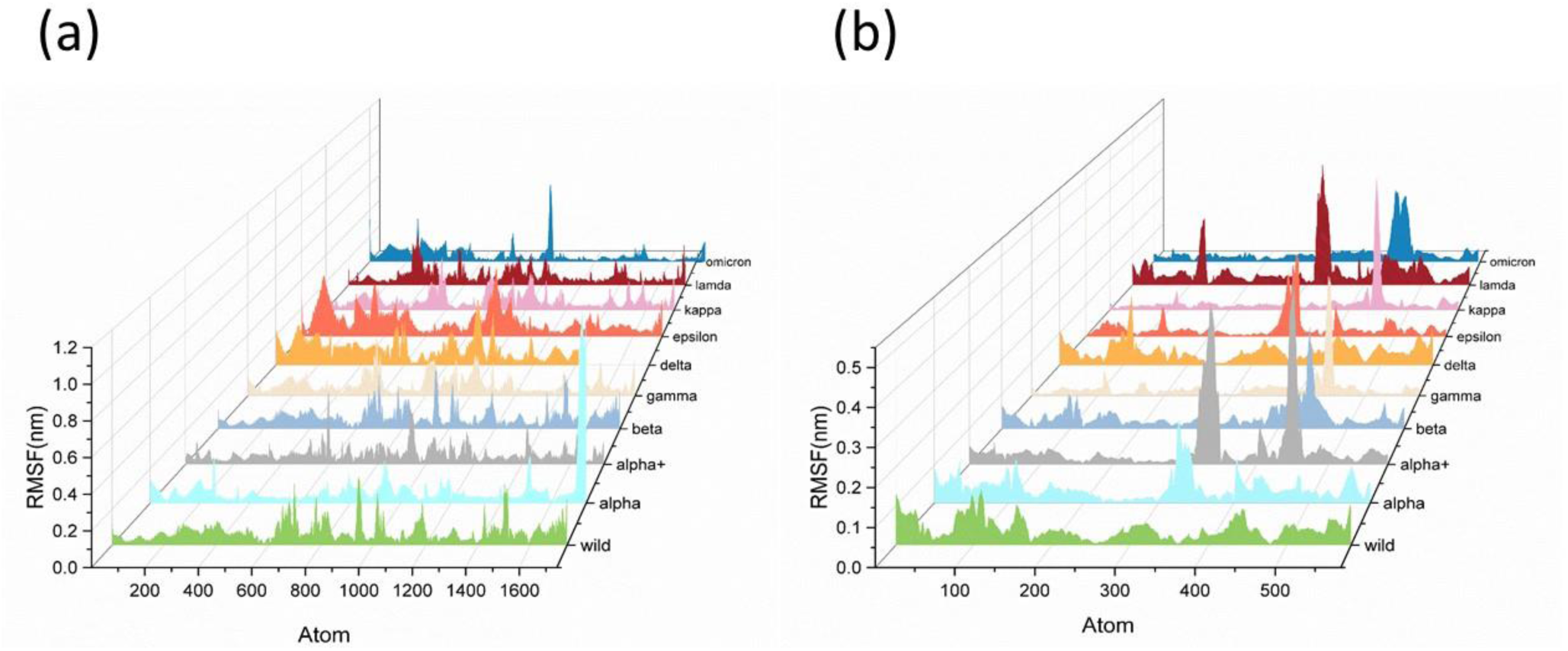
The root means square fluctuations of backbone atoms obtained from PCA (a) ACE2 receptor and (b) RBD wild and mutants. The PCA-based RMSF values show essential fluctuations, which is corresponding to the residues that are associated with secondary structural changes as well as the higher fluctuations at binding and mutant sites.

**Figure S5.**
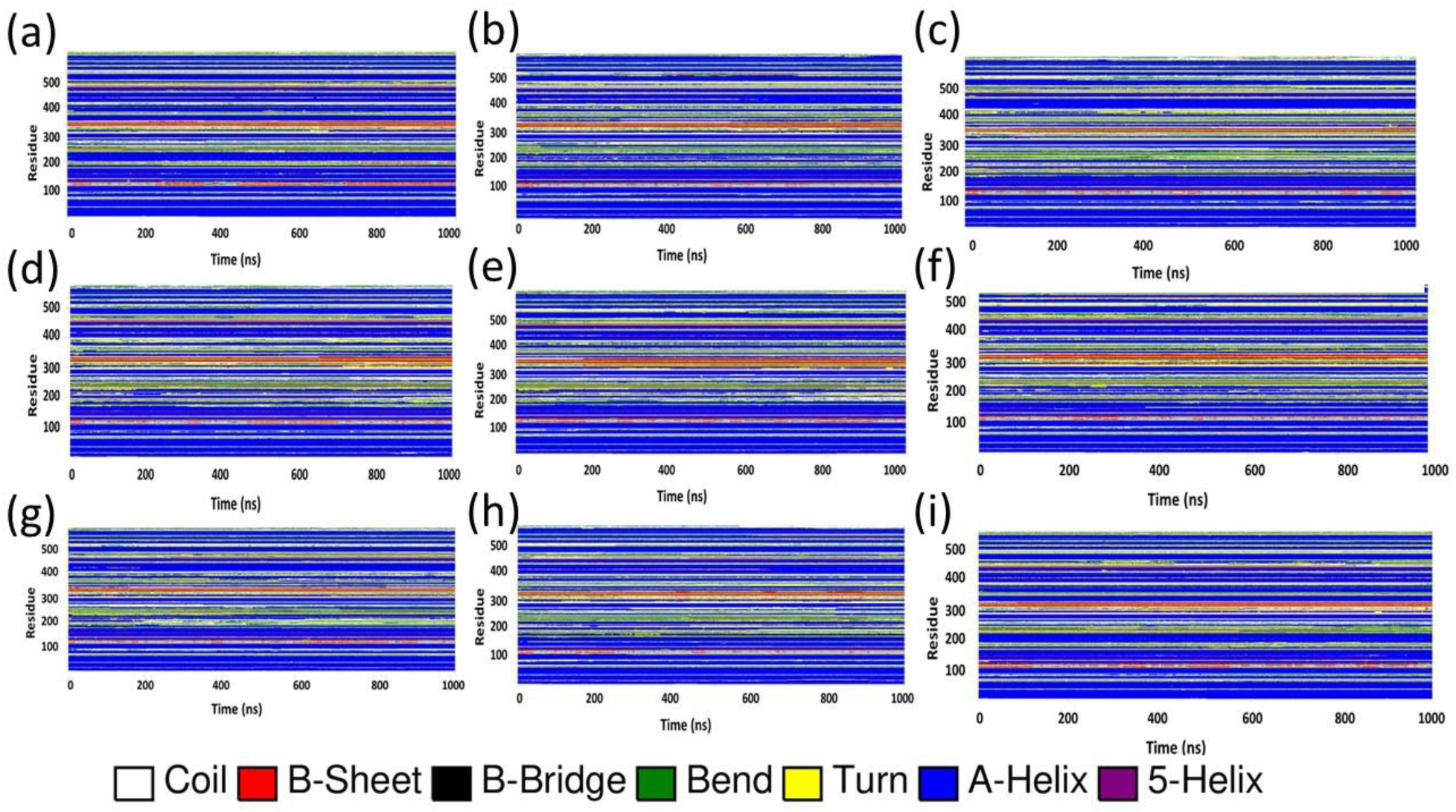
Time-dependent secondary structural changes of the ACE2 receptor. (a) wild,(b)alpha,(c)beta,(d)gamma,(e)delta,(f)epsilon,(g)kappa,(h)lambda, and (i) omicron.

**Figure S6.**
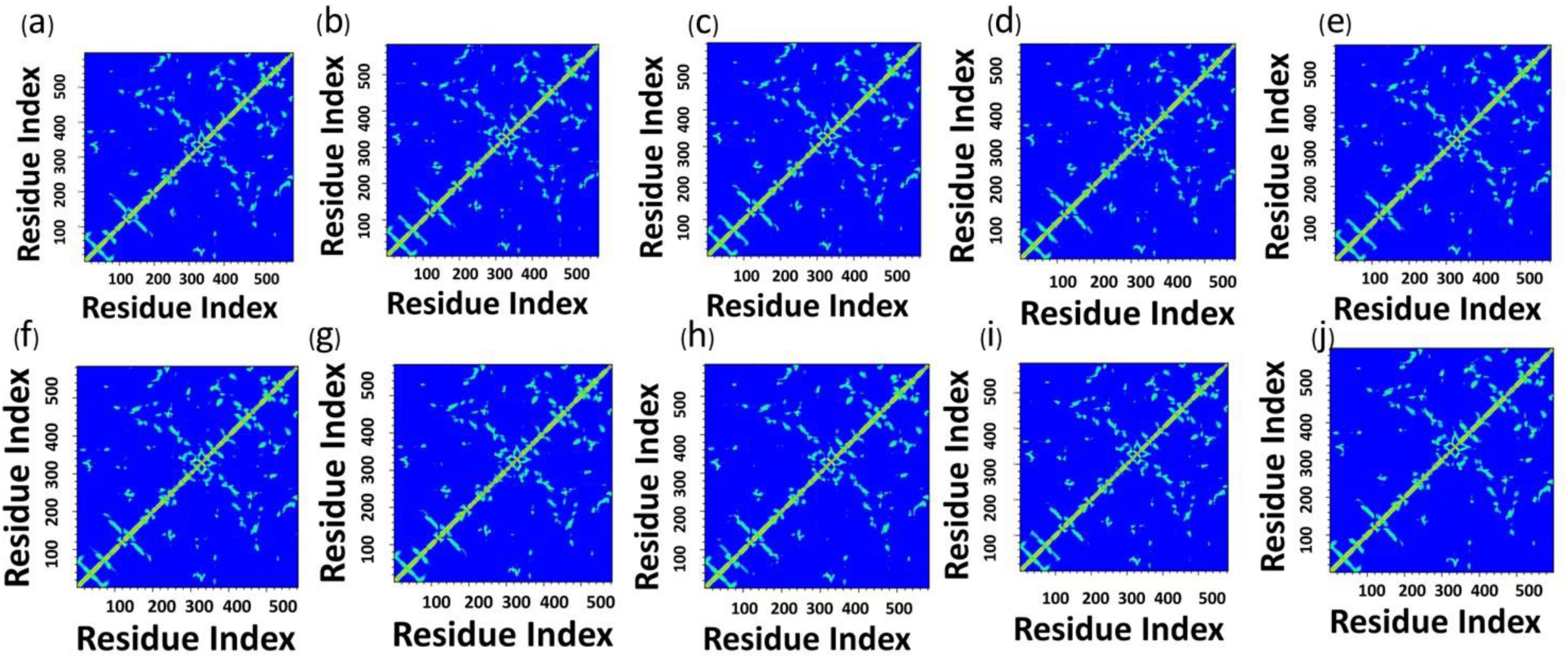
The residue-residue distance matrix was constructed for the ACE2 receptor. (a) Wild, (b) alpha, (c) alpha+, (d) beta, (e)gamma,(f)delta,(g)epsilon,(h)kappa,(i)lambda, and (j) omicron.

**Figure S7.**
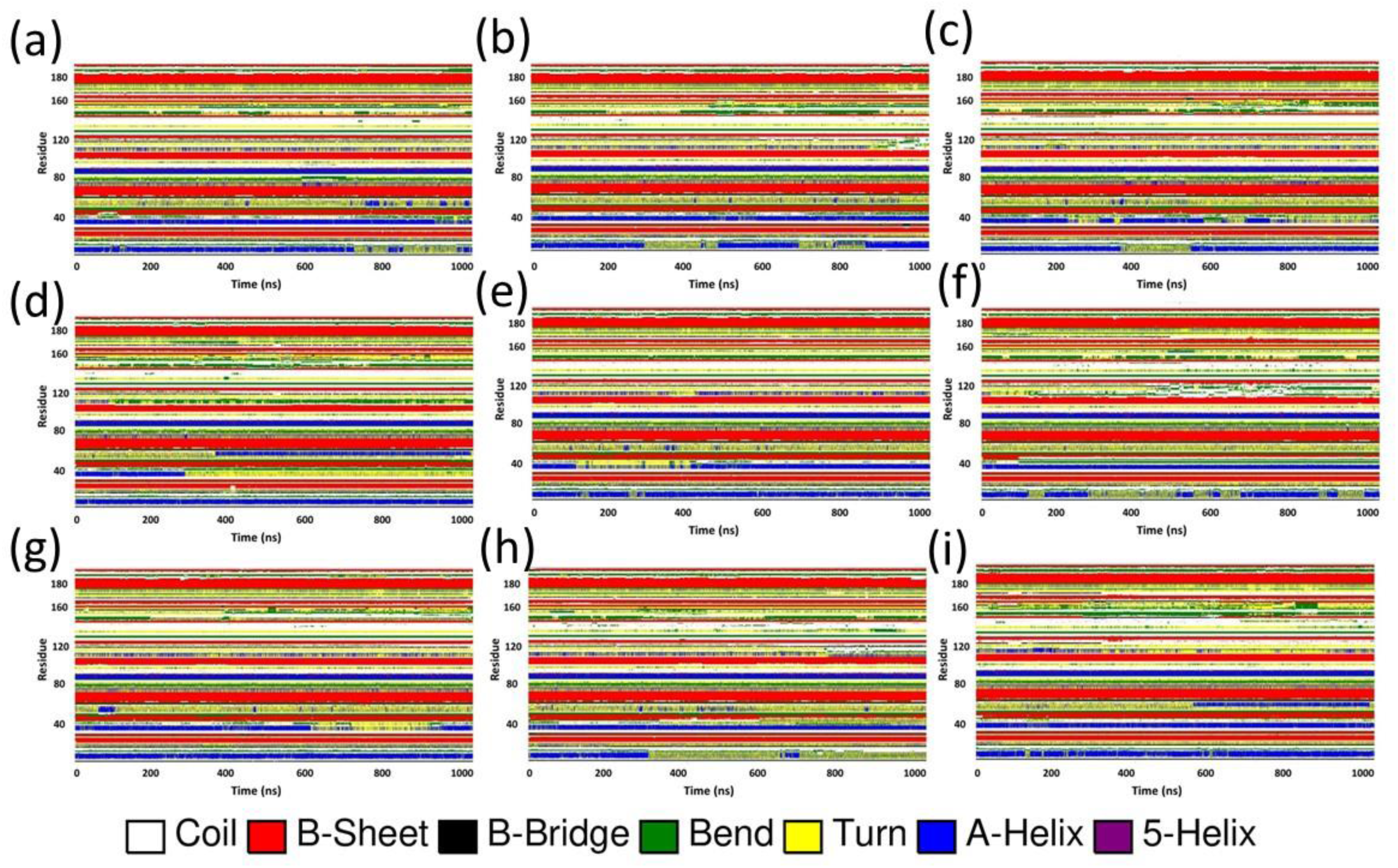
Time-dependent secondary structural changes of the RBD. (a) wild,(b)alpha,(c)beta,(d)gamma,(e)delta,(f)epsilon,(g)kappa,(h)lambda, and (i) omicron.

**Figure S8.**
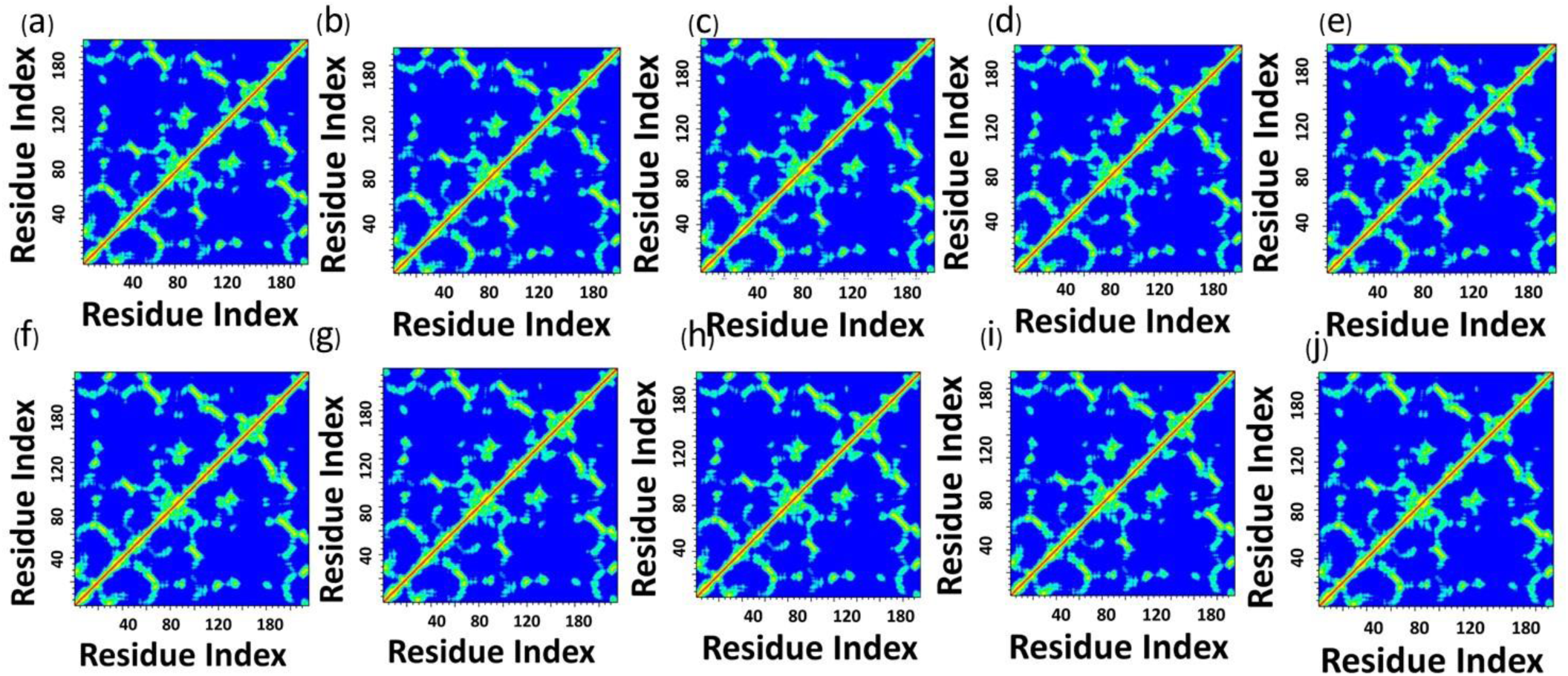
The residue-residue distance matrix was constructed for RBD. (a) Wild, (b) alpha, (c) alpha+, (d) beta, (e)gamma,(f)delta,(g)epsilon,(h)kappa,(i)lambda, and (j) omicron.

**Figure S9.**
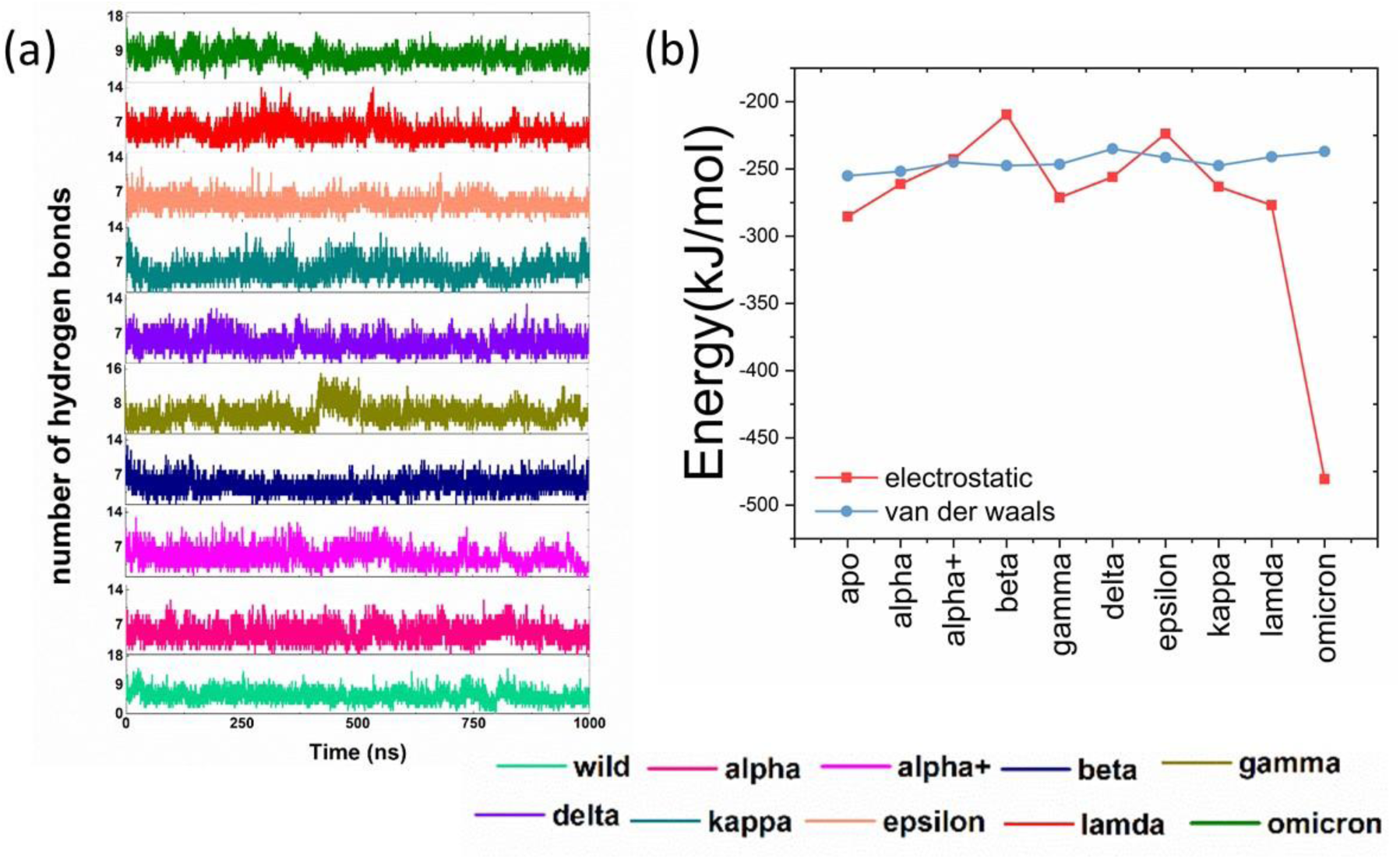
The number of hydrogen bonds formed between ACE2-RBD and mutants (a) and the average electrostatic and van der Waals interaction energy between ACE2-RBD mutants.

**Figure S10.**
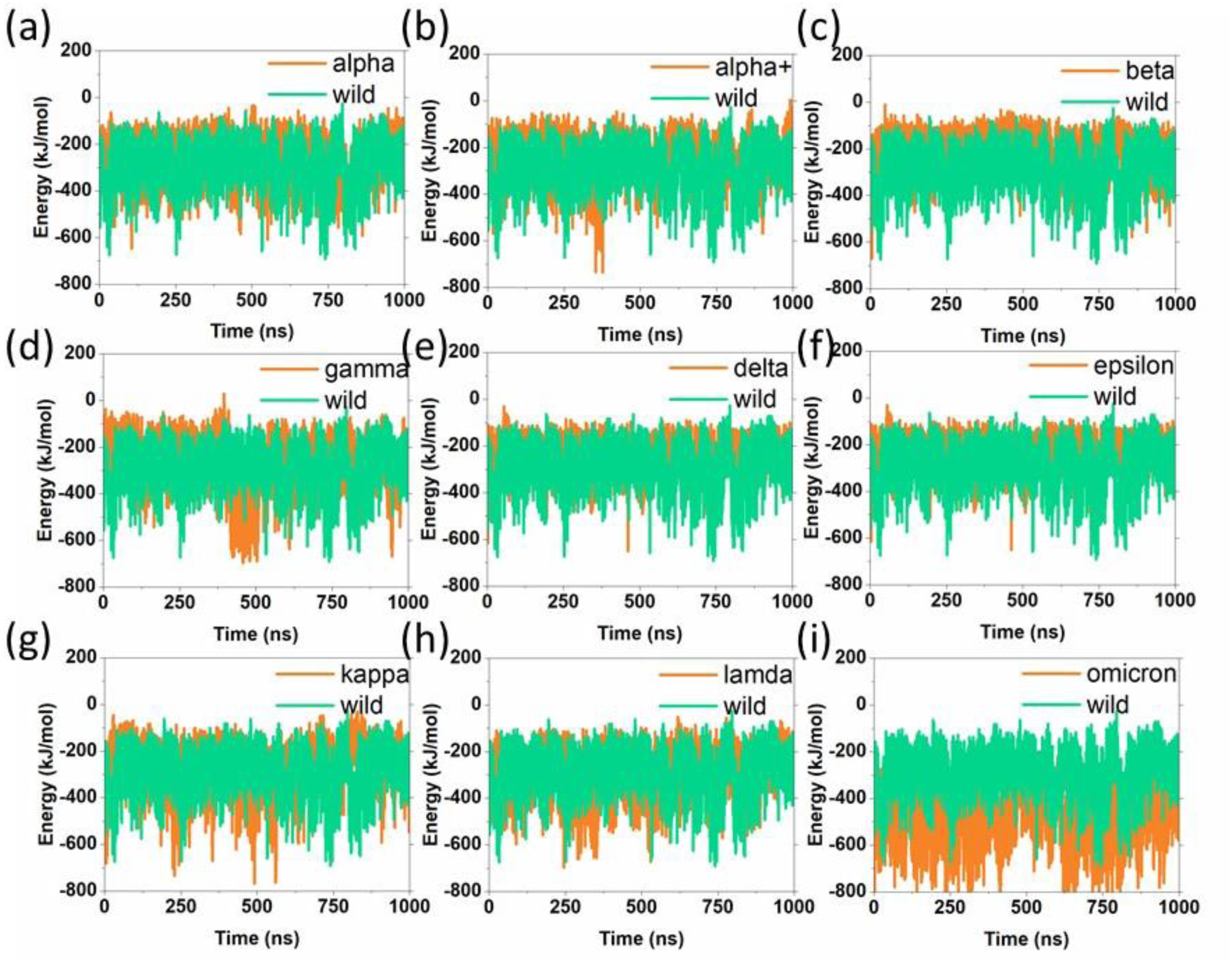
The electrostatic interaction energy over time between ACE2-RBD and mutants.

**Figure S11.**
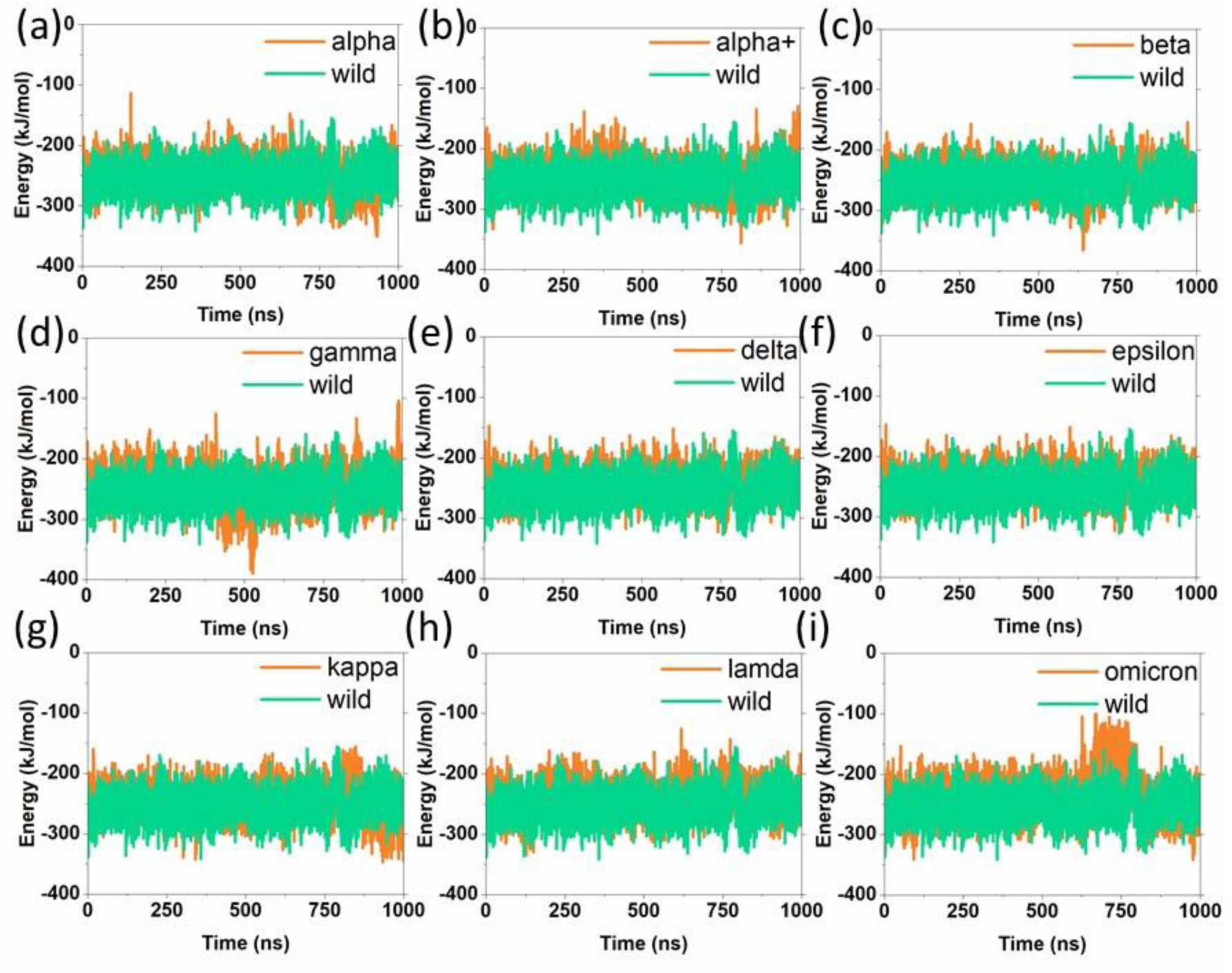
The van der Waals interaction energy over time between ACE2-RBD and mutants.

**Figure S12.**
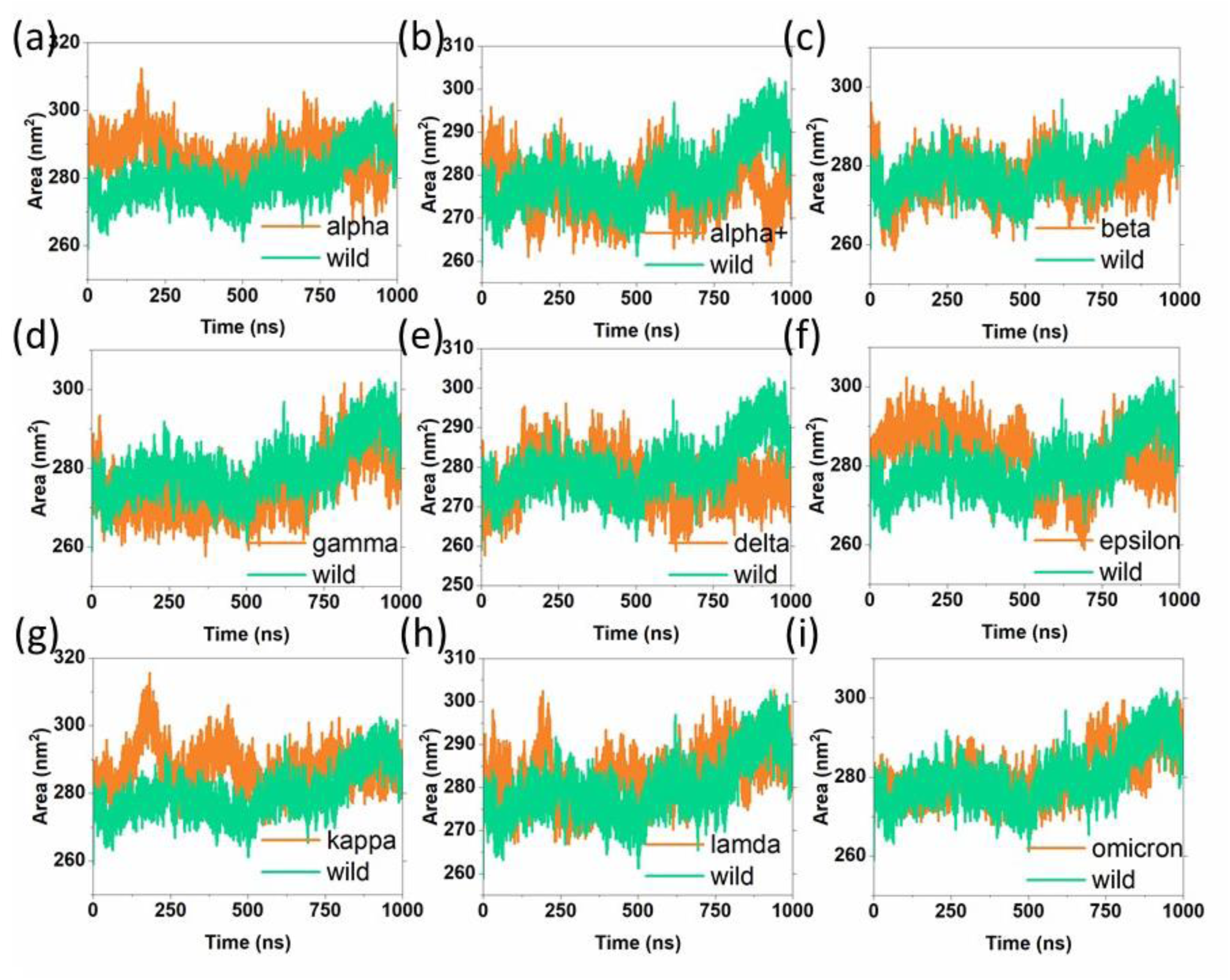
The time-dependent solvent accessible surface area for ACE2 receptors.

**Figure S13.**
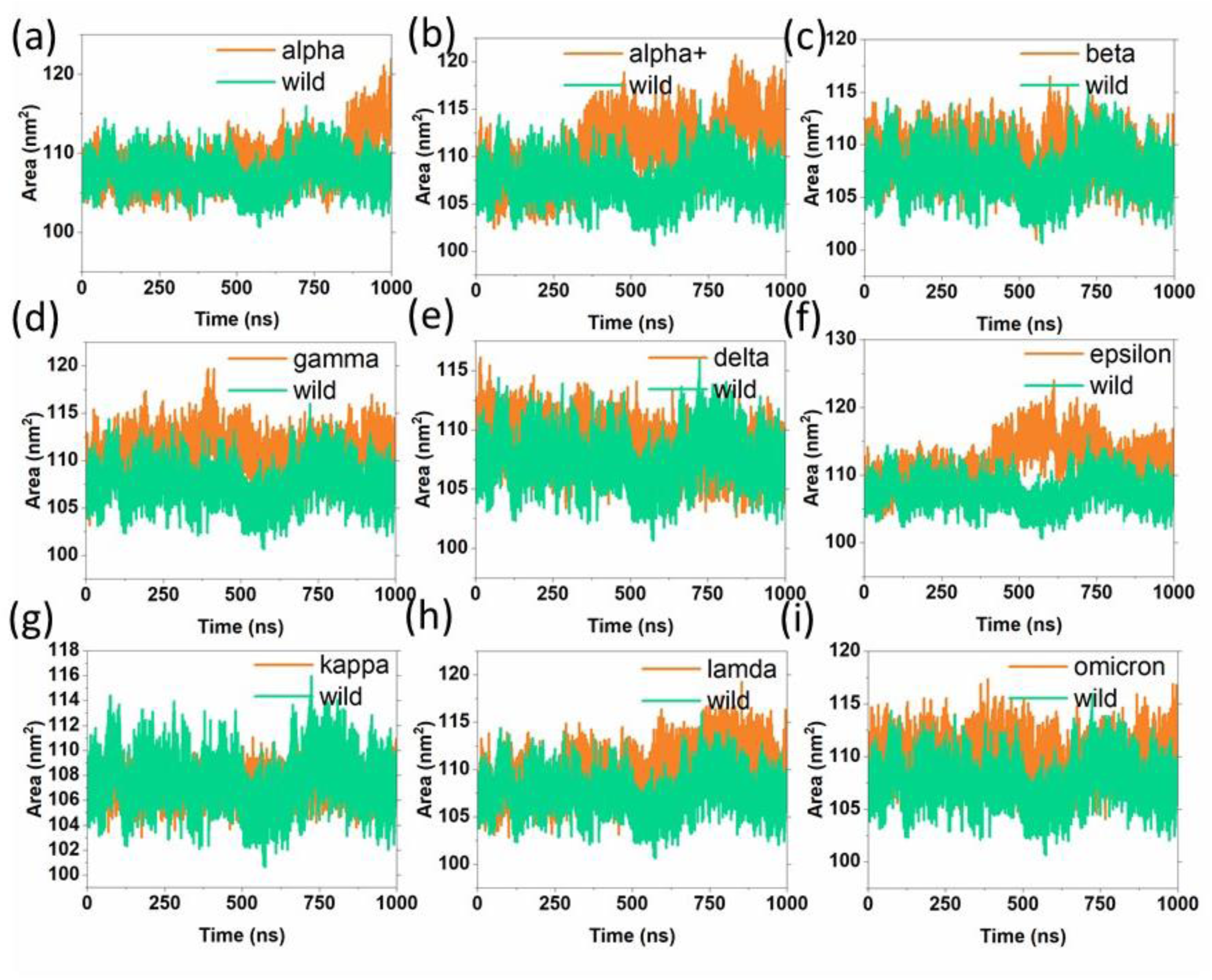
The time-dependent solvent accessible surface area of RBD wild and its mutants.

**Figure S14.**
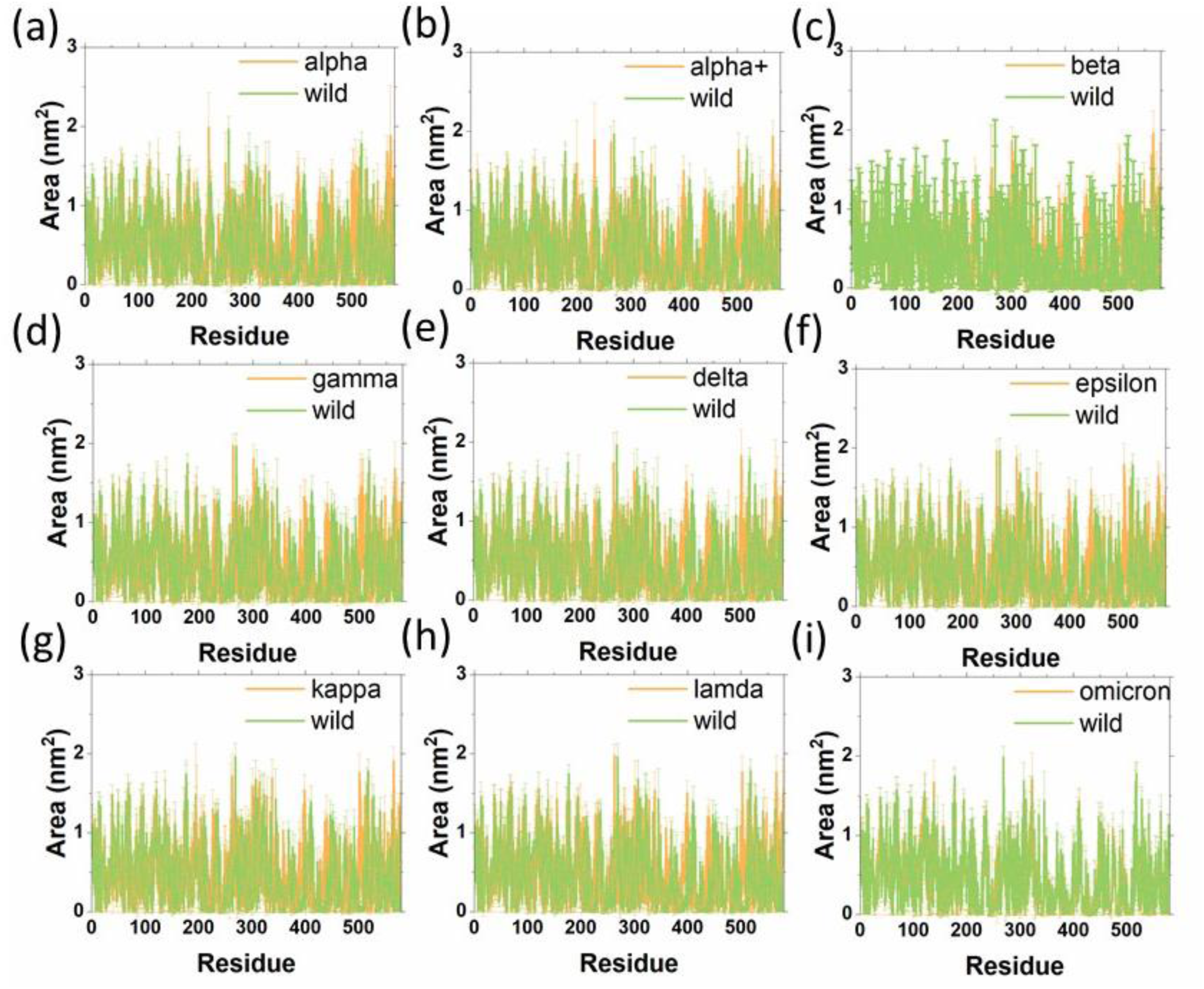
The residue-wise solvent accessible surface area of ACE2 receptors.

**Figure S15.**
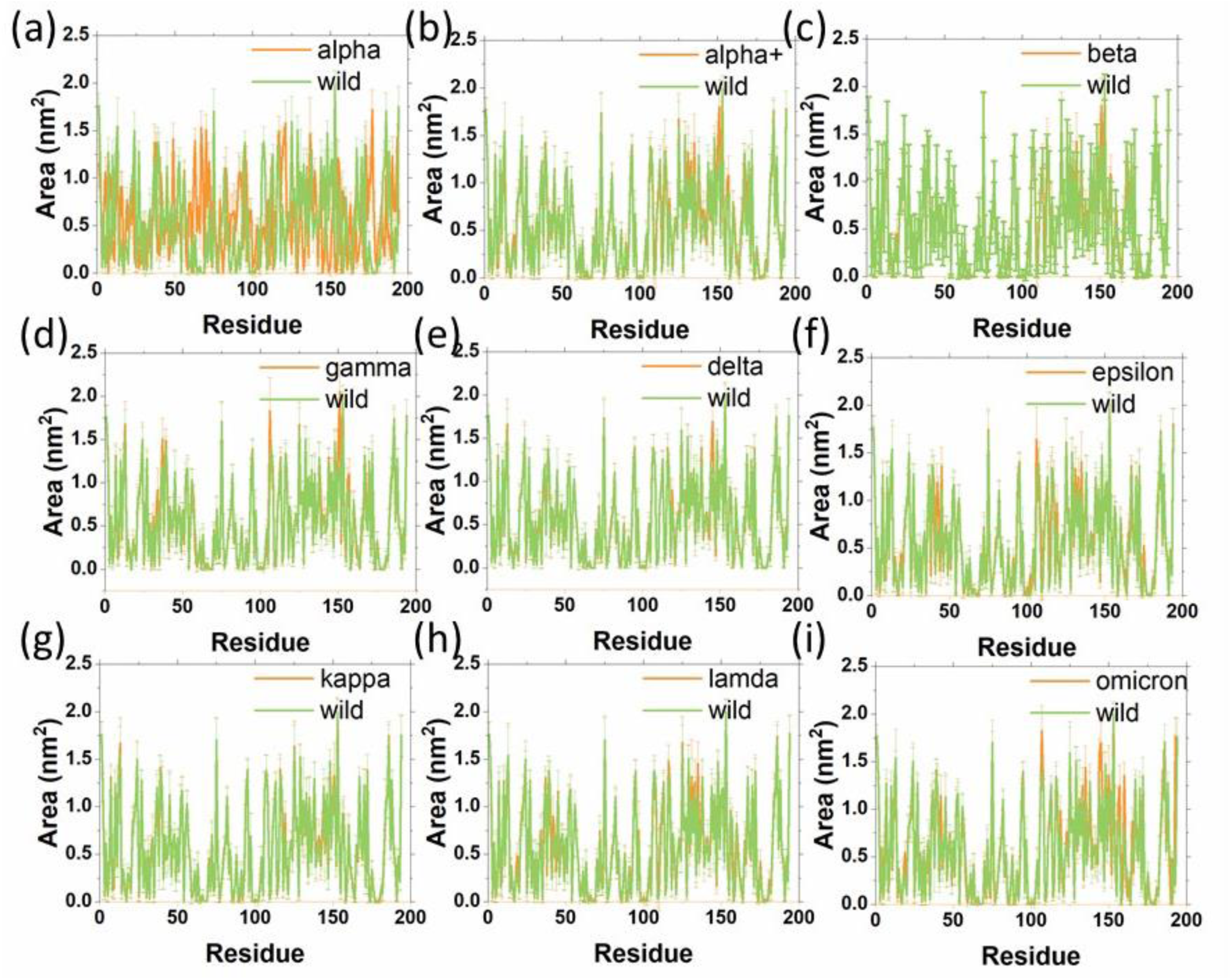
The residue-wise solvent accessible surface area of RBD and its mutants.

**Figure S16.**
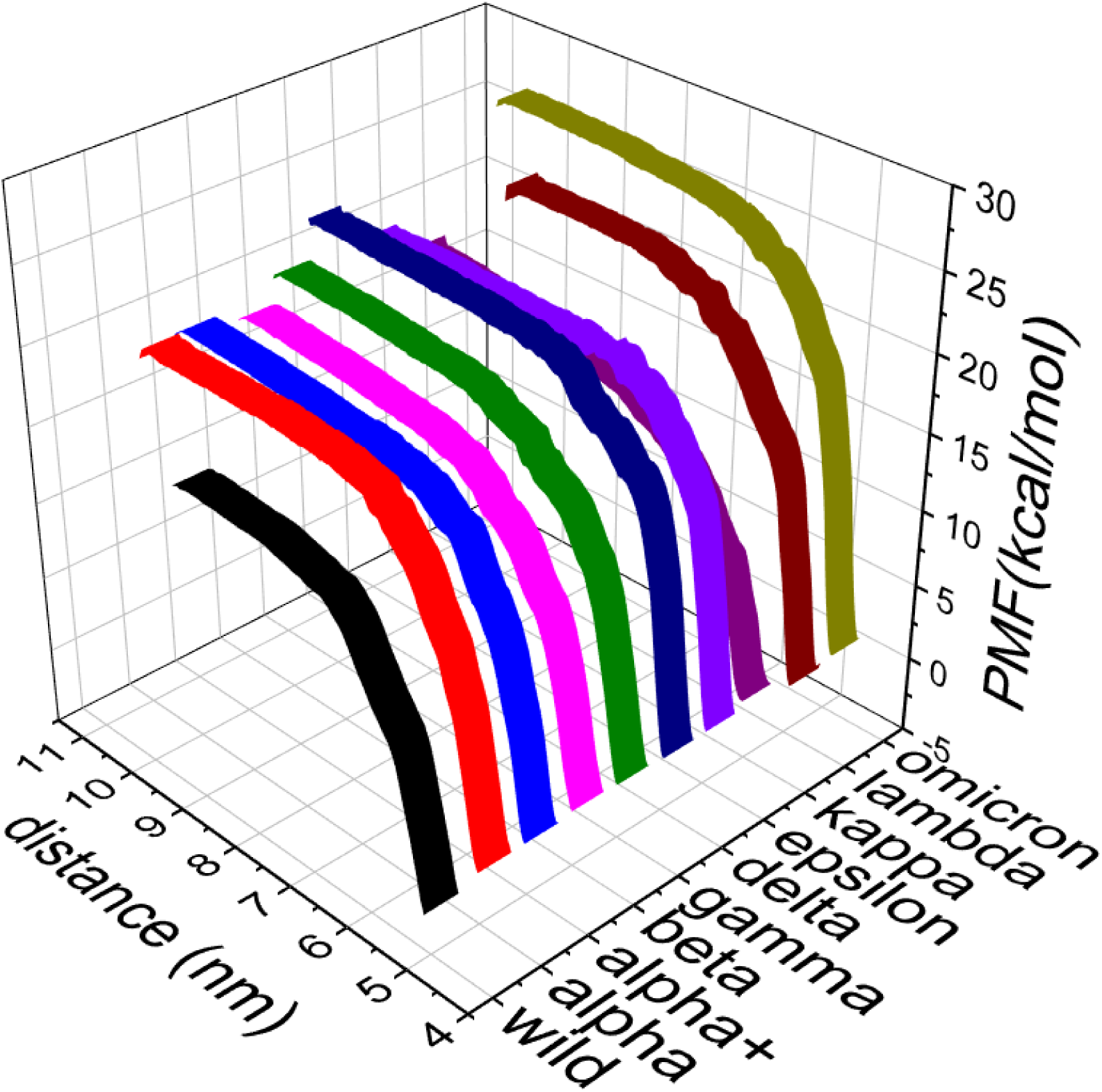
The potential of mean force of ACE2-RBD wild and mutants. The omicron variant shows the highest binding energy compared to wild and other mutants.

**Figure S17.**
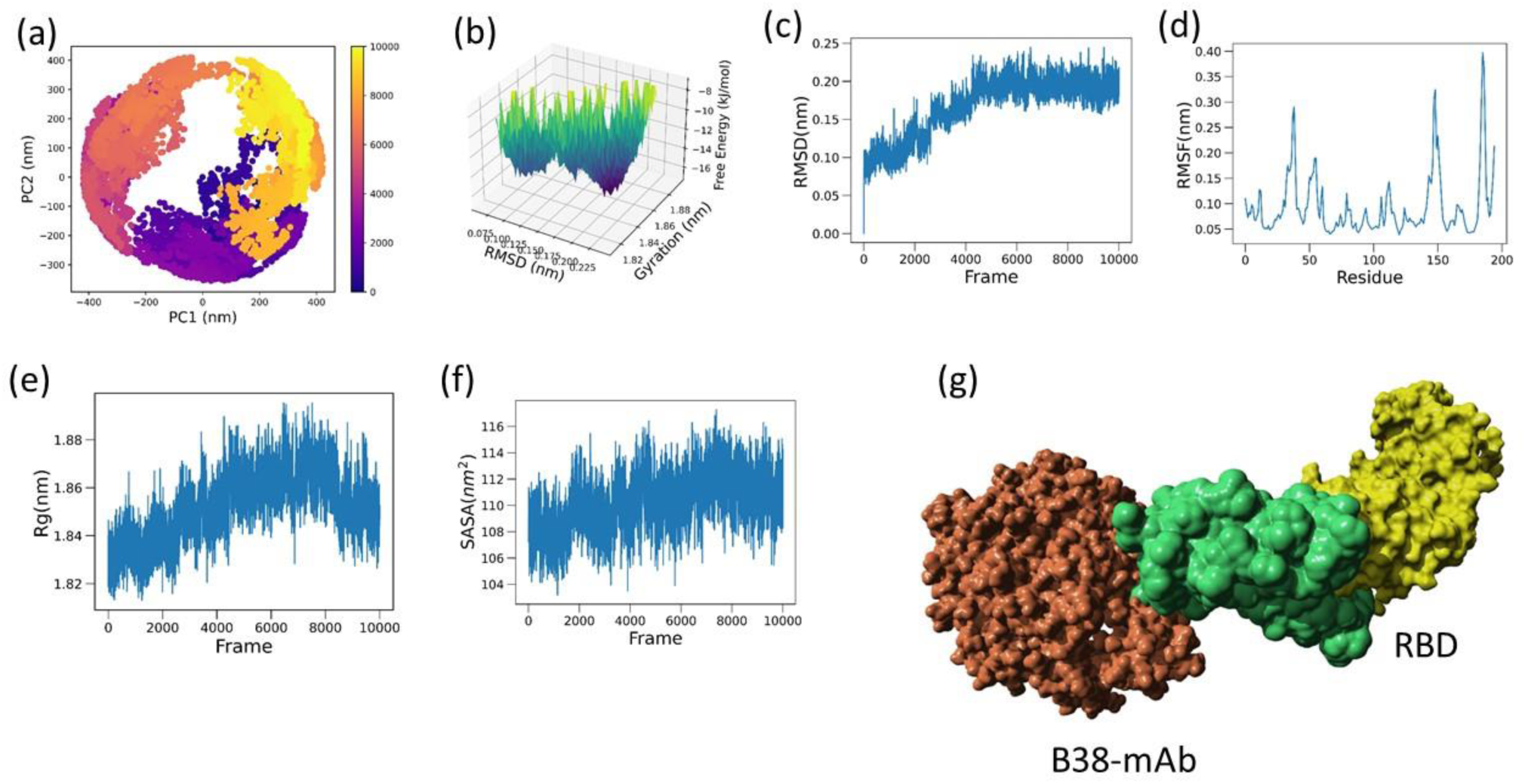
(a) Represents the PCA of RBD that is bound with antibody and (b) represents the free energy landscape of RBD obtained from RMSD of backbone atoms (c) and radius of gyration (e). The RMSF of Cα atoms and 3D structural representation of B38 antibody-bound RBD are given in (f) and (g) respectively.

**Figure S18.**
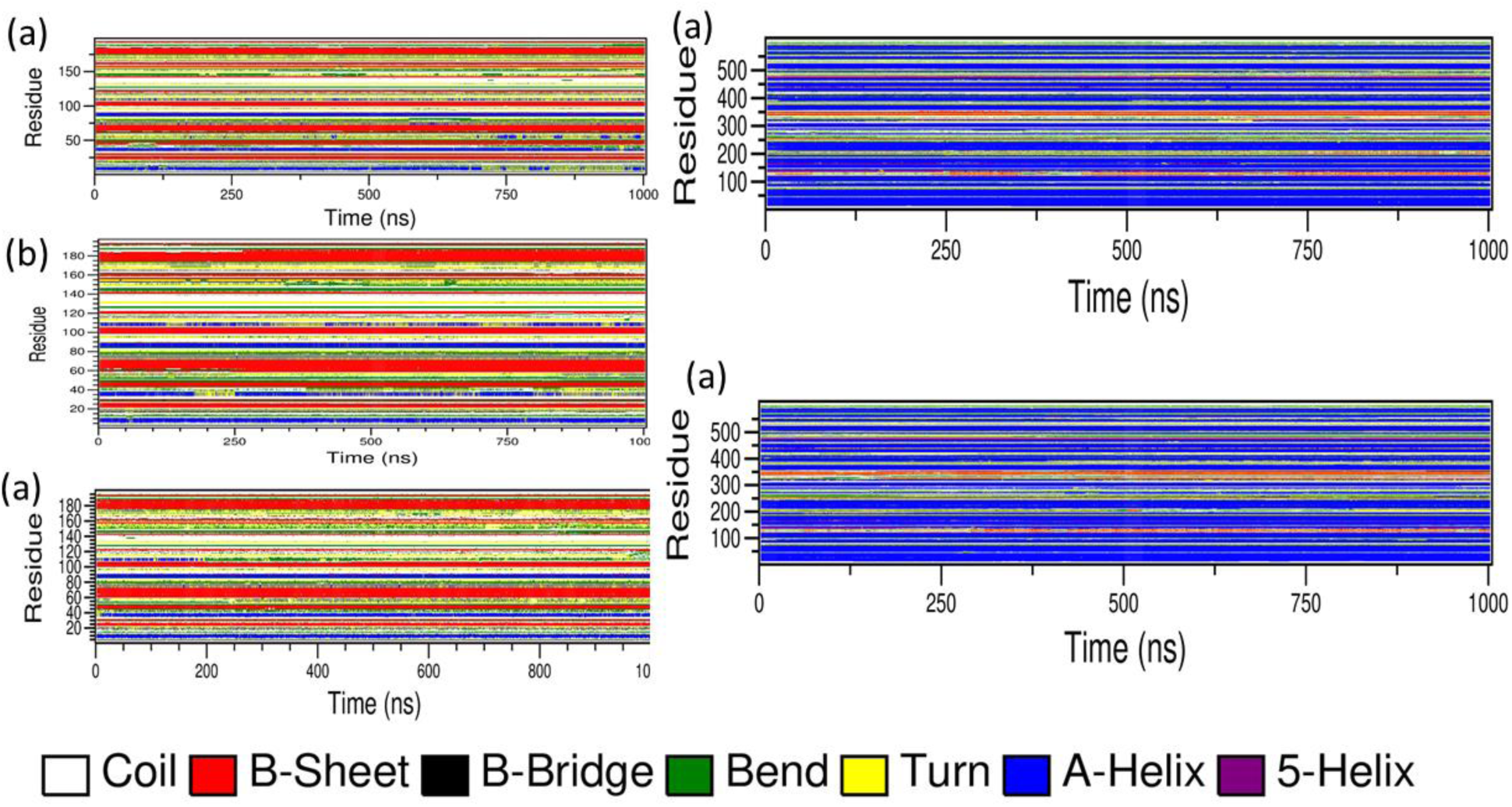
The time-dependent secondary structural changes of RBD (a) wild (b) antibody bound and (c) ACE2 bound RBD that inhibited by the antibody. (d) and (e) represents the time defendant secondary structural changes of wild ACE2 receptor and that bound with inhibited RBD.

**Table S1.**
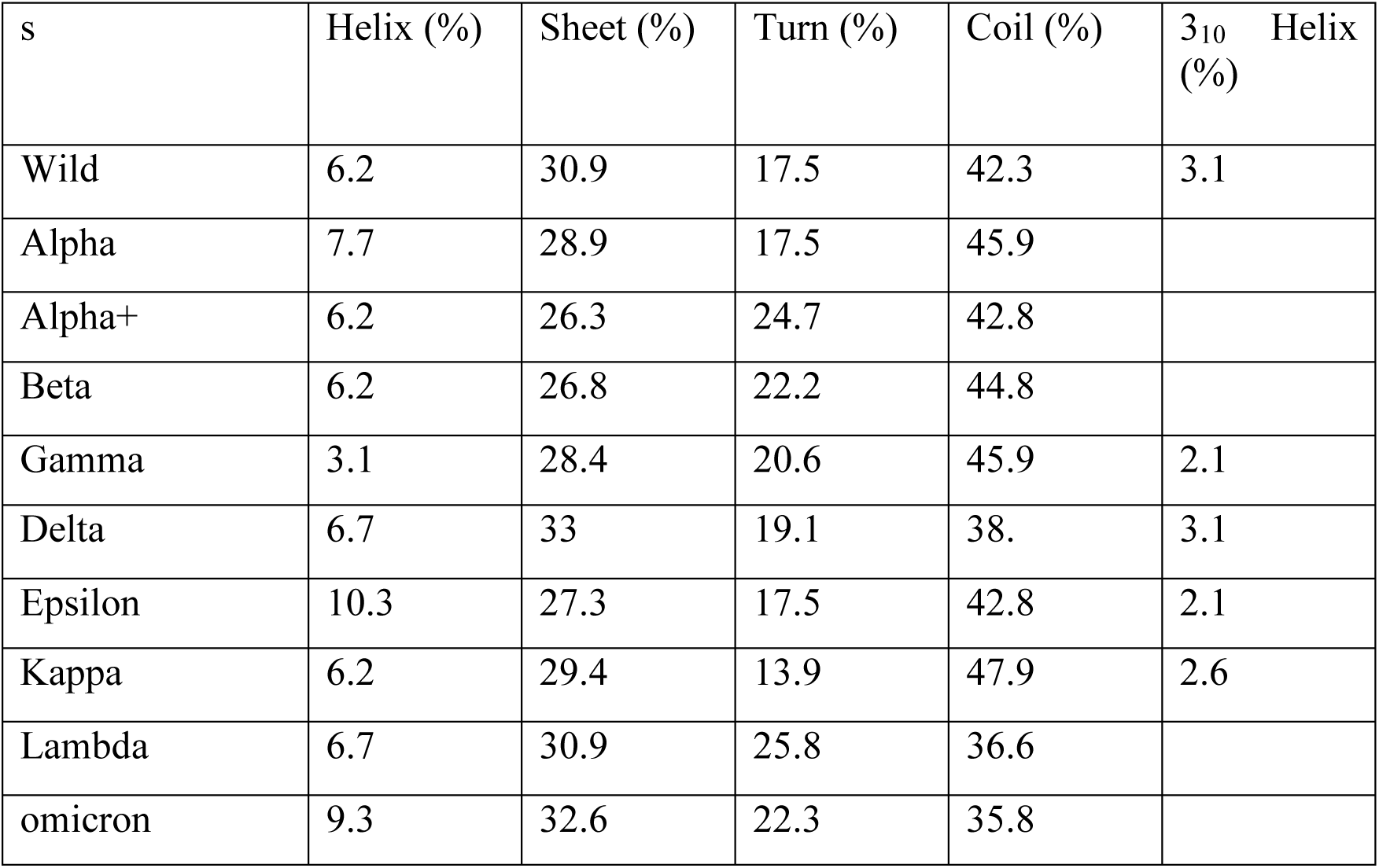
The percentage of the secondary structure after 1 µs simulations of ACE2 receptor.

## Notes

### Competing Interest Statement

The authors have declared no competing interest.

### Summary of Updates

The revised version, is updated with compuational details and high resolution figures.

